# Structured recurrence of latent neural processes predicts memory recognition performance

**DOI:** 10.64898/2026.10.16.759557

**Authors:** K. F. Christensen, M. Klarlund, G. Fernández-Rubio, M. Rosso, S. Streton, M. Schiassi, M. L. Kringelbach, P. Vuust, L. Bonetti

## Abstract

Understanding cognition requires explaining not only which neural processes are engaged, but how their relationships evolve through time. We tested whether the recurrent organization of low-dimensional whole-brain dynamics relates to auditory recognition behaviour beyond activation magnitude. After a brief learning phase, 65 participants underwent magnetoencephalography (MEG) while recognizing three-, five-, and seven-sound auditory sequences, presented unchanged or with a single early or late tone alteration. Broadband Brain Network Estimation via Source Separation (BROAD-NESS) identified broadband latent networks from source-reconstructed activity and represented their joint activation as trajectories through a multidimensional state space; recurrence quantification analysis characterized their temporal organization. Memorized melodies elicited more recurrent states and longer, more deterministic and persistent recurrence structures, together with lower divergence, than changed melodies. Across participants, structured recurrence was consistently associated with higher accuracy and faster responses, whereas recurrence frequency showed no consistent behavioural association. After adjustment for condition, melody length, their interaction, and matched root-mean-square activation amplitude, recurrence measures increased marginal *R*^2^ by 0.273. Nested participant-wise cross-validation reduced held-out prediction error by 11.81% (95% CI: 4.01%–17.65%). These findings support a view of cognition in which performance depends critically on how low-dimensional neural processes jointly organize and evolve over time, beyond how strongly they are activated.

## Introduction

A central challenge in cognitive neuroscience is to explain how distributed neural activity gives rise to behaviour. Neural processes unfold continuously and collectively, yet neural–behaviour relationships are commonly studied through the magnitude or timing of responses measured in individual regions, components, or frequency bands ^1–14^. These measures reveal when and how strongly particular processes are engaged, but they provide limited information about how distributed processes evolve together. If cognition emerges from coordinated neural dynamics, identifying which processes are active is only part of the explanation, and their joint temporal organization may be equally or even more important.

The low-dimensional hypothesis for cognition formalizes this idea through two linked principles ^15^. First, coordinated whole-brain activity gives rise to a limited number of latent processes that can be recruited and reconfigured according to internal and external demands. Second, cognition depends on the interactions and changing relationships among these processes. Cognitive richness may therefore arise not from an unlimited number of independent neural dimensions, but from the ways in which a smaller set of processes combine, constrain one another, and reorganize over time. This account makes a specific prediction: cognitive performance should reflect not only the presence or activation strength of latent processes, but also the temporal organization produced by their joint evolution.

Low-dimensional organization has emerged as a fundamental property of complex dynamical systems, whose apparently high-dimensional behaviour can often be captured by a much smaller set of interacting degrees of freedom ^16^. In the brain, neural-manifold studies reveal low-dimensional trajectories, hidden-state analyses reveal recurring configurations and transitions between brain states, and dynamic-connectivity and latent-network models reveal reconfiguration at rest and during cognitive tasks ^17–25^. Together, these findings indicate that brain activity is organized through a relatively small set of latent processes whose configurations and dynamics vary over time. What remains unresolved is what about this organization specifically matters for cognition and how it translates into behaviour.

To examine how this question has been addressed in studies directly relating neural activity to behavioural performance, we conducted a targeted screen of 59 recent human electrophysiological studies of memory, recognition, and sequence learning. Of these, 71% used evoked-response magnitude or timing or oscillatory magnitude as their behavioural neural measure (Table S1; individual study classifications are reported in Table S2). Evoked-response amplitudes and latencies have been related to auditory working-memory capacity, successful sound and melody recognition, and recognition precision^1,2,4,5^, while oscillatory power has been associated with auditory-memory accuracy, response speed, and benefits from selective attention3^,6,8–10^. Thus, whereas studies of low-dimensional neural dynamics have characterized how distributed neural processes are organized and reconfigured, studies directly relating neural activity to behaviour have predominantly focused on response strength and timing. These findings establish that response strength and timing are behaviourally relevant but leave unresolved whether cognitive performance is also supported by how distributed neural processes organize and evolve together over time. If their joint dynamics are consequential for cognition, this organization should explain individual cognitive performance beyond the activation strength of the latent processes themselves.

Auditory sequence recognition provides a particularly stringent test of this possibility because successful recognition depends inherently on temporal integration: each sound must be interpreted in relation to what preceded it, compared with a learned representation, and used to constrain what should follow. MEG studies have traced how such sequences are encoded, recognized, reactivated, and mentally manipulated across time ^26–31^. Related work has shown that recognition and prediction errors recruit distributed auditory, cingulate, medial temporal, and prefrontal systems whose contributions evolve across the sequence^32–37^. Combined with the temporal resolution of MEG, this paradigm makes it possible to follow whole-brain trajectories as recognition unfolds and to test whether their temporal organization is related to behavioural performance.

To this aim, participants evaluated three-, five-, and seven-tone excerpts extracted from a previously memorized musical piece, presented unchanged or with a single early or late tone alteration, while MEG was recorded. We analysed the source-reconstructed activity with the Broadband Brain Network Estimation via Source Separation (BROAD-NESS) toolbox^38^, developed in the context of the Network Estimation via Source Separation (NESS) framework^39–41^, an integrated platform that identifies broadband latent networks and represents their simultaneous activation as trajectories through a multidimensional state space. Within this framework, recurrence quantification analysis (RQA) characterized the returns, repeated sequences, persistence, and divergence of the trajectories. We asked whether recurrent organization differed between memorized and changed melodies, whether individual variation in this organization was associated with recognition accuracy and response speed, and whether it explained behaviour beyond the mere activation of the same latent processes. We predicted that memorized melodies would elicit more stable and structured trajectories, that stronger diagonal and vertical recurrence structure would accompany better performance, and that greater trajectory divergence would accompany poorer performance.

## Methods

### Participants

To increase the sample while broadening its cultural representation, we recruited 65 participants in Denmark: 34 Danish participants and 31 recently arrived Chinese immigrants. The sample comprised 41 female and 24 male participants. Mean age was 24.4 years (SD = 4.2; range 19–41). All participants were healthy and reported normal hearing (see Table 1).

**Table 1:**
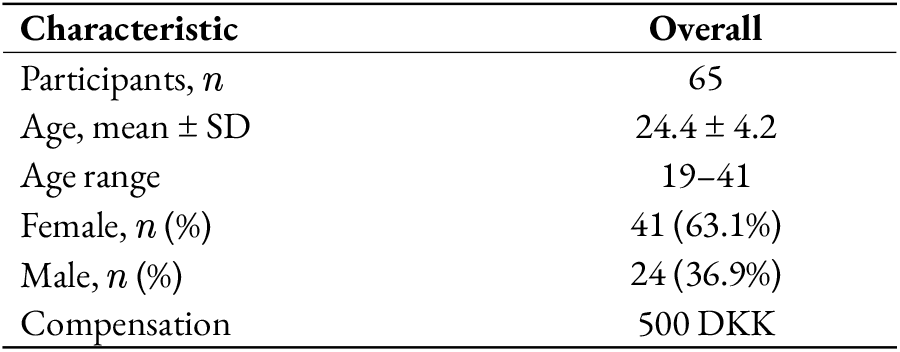
Participant characteristics. Percentages use the full sample as denominator.

| Characteristic | Overall |
| --- | --- |
| Participants, $n$ | 65 |
| Age, mean ± SD | 24.4 ± 4.2 |
| Age range | 19–41 |
| Female, $n$ (%) | 41 (63.1%) |
| Male, $n$ (%) | 24 (36.9%) |
| Compensation | 500 DKK |

The study was approved by the Institutional Review Board (IRB) of Aarhus University (case number: DNC-IRB-2023-009) and conducted in accordance with the Declaration of Helsinki – Ethical Principles for Medical Research. All participants provided informed consent prior to the experiment and received 500 DKK for their participation.

### Data Inclusion and Missingness

Some melody-length blocks were excluded during MaxFilter processing because of data-quality problems. The retained samples therefore comprised 61 participants for three-tone melodies (M3), 60 for five-tone melodies (M5), and 62 for seven-tone melodies (M7). These block-specific sample sizes should not be interpreted as successive stages of participant exclusion. 12 participants were missing only one of the three melody-length blocks: 4 lacked M3, 5 lacked M5, and 3 lacked M7. Each of these participants still contributed data to analyses of the other two melody lengths. Consequently, analyses conducted separately by melody length used all 60–62 available participants, whereas cross-melody repeated-measures analyses were restricted to the 53 participants represented at all three melody lengths. Thus, the reduction to 53 reflects the requirement for a complete set of three blocks within each participant, rather than the loss of all data from those participants.

Trial-level quality exclusions were shared across the behavioural and neural analyses: 152 trials overlapping recorded bad-data segments and three trials lacking a corresponding MEG epoch were removed. Behavioural summaries retained correct, incorrect, and no-response trials after these exclusions. The condition-specific neural averages supplied to BROAD-NESS used only correct-response trials from the same retained set.

No participant had a recurrence rate of zero in any analysed condition. The RQA samples therefore matched the retained block samples: 61 participants for M3, 60 for M5, and 62 for M7, representing 65 unique participants. Cross-melody RQA analyses included the 53 participants with all three blocks.

### Experimental Design

Participants completed an auditory recognition task during MEG recording. They first listened twice to the first four bars of the right-hand part of Johann Sebastian Bach’s Prelude No. 2 in C Minor, BWV 847, and were instructed to memorize it. A 20-s rest period followed the learning phase. During recognition, participants heard excerpts containing three (M3), five (M5), or seven tones (M7) and judged whether each excerpt had been taken unchanged from the memorized piece (memorized; M) or contained a change.

Three memorized excerpts were used at each melody length and were each presented 12 times, giving 36 memorized trials per block. Changed excerpts were derived from the same musical material by replacing a single tone. The replacement occurred either relatively early or at the end of the excerpt: tone 2 or 3 in M3, tone 3 or 5 in M5, and tone 5 or 7 in M7, see Figure 1 for a graphical representation of this. Variants of the replacement pitch served as stimulus exemplars and were pooled within change position. Each melody-length block therefore comprised 72 trials: 36 memorized, 18 early-change, and 18 late-change trials. Consecutive tones had an onset-to-onset interval of 350 ms.

**Figure 1:**
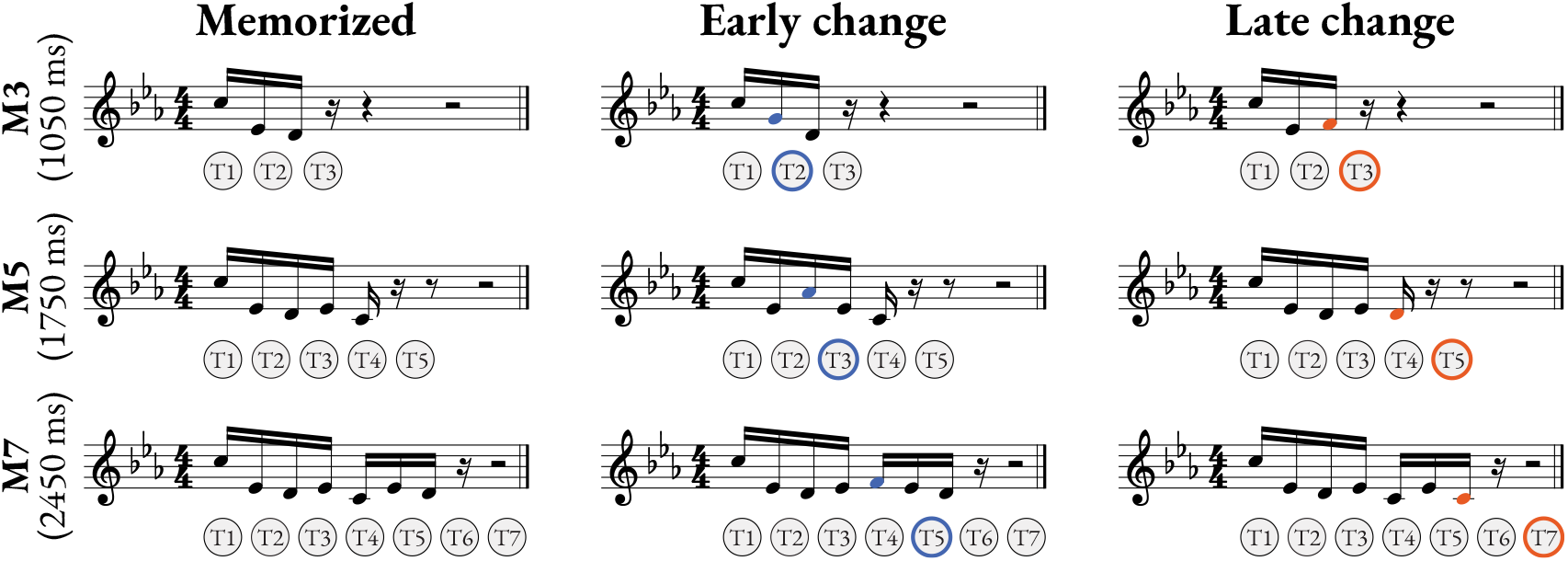
Example musical notation and tone schematics illustrate memorized melodies of three, five, and seven tones and the single-tone alterations in 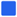 early-change and 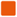 late-change trials.

The order of the three melody-length blocks was randomized across participants, and all memorized and changed trials were randomized together within each block. Before each block, participants were informed of the number of tones in the upcoming excerpts. They responded using one button for memorized and another for changed. Responses were accepted for 3.0 s in M3, 3.7 s in M5, and 4.5 s in M7, measured from excerpt onset. A trial on which no response was registered before the corresponding deadline was classified as a no-response trial. The task was presented using PsychoPy, and stimulus-onset triggers were sent to the MEG acquisition system.

### Behavioural Analysis

Responses were classified as correct when a memorized excerpt received a memorized response or a changed excerpt received a changed response. All other registered responses were classified as incorrect, while trials reaching the response deadline were classified as no responses. After applying the shared trial-quality mask, response proportions were calculated separately for every combination of participant, melody length, and condition. Reaction times were averaged separately for correct and incorrect responses within each melody-length and condition combination; no-response trials did not contribute to reaction-time means. A reaction-time mean was treated as missing when a participant made no registered response for that combination.

The proportions of correct, incorrect, and no responses were each submitted to a 3 × 3 repeated-measures analysis of variance (ANOVA), with condition (memorized, early change, and late change) and melody length (M3, M5, and M7) as within-participant factors. The same factorial design was applied separately to mean reaction times for correct and incorrect responses. These omnibus tests assessed the main effects of condition and melody length and their interaction. Omnibus *p*-values were FDR-corrected across the three effects within each response category, and effect sizes were expressed as partial *η*^2^.

Comparisons displayed by significance brackets in the behavioural figures were based on paired-samples *t*-tests of participant-level values. Only participants with finite values in both conditions contributed to a given reaction-time comparison. The resulting *p*-values were adjusted using FDR correction across the family of brackets displayed in the relevant figure. Adjusted significance was denoted by * for *p* < .05, ** for *p* < .01, and *** for *p* < .001.

### Neural Data Acquisition

#### MEG Acquisition

MEG was recorded in a magnetically shielded room at Aarhus University Hospital using an Elekta Neuromag TRIUX system with 306 channels (204 planar gradiometers and 102 magnetometers). Data were acquired at 1000 Hz with an analogue passband of 0.1–330 Hz. Before recording, the participant’s head shape and the locations of four head-position-indicator coils were registered relative to three anatomical landmarks using a Polhemus Fastrak 3D digitizer. The coils tracked head position continuously during acquisition and enabled subsequent movement correction. Bipolar electrooculography and electrocar-diography channels recorded eye movements, blinks, and cardiac activity for artefact identification.

#### MRI Acquisition

Structural MRI was acquired on a CE-approved 3 T Siemens scanner at Aarhus University Hospital on a separate day from MEG. T1-weighted images were obtained using a fat-saturated MPRAGE sequence with 1 mm isotropic resolution, echo time 2.61 ms, repetition time 2300 ms, a reconstructed matrix of 256 × 256, echo spacing 7.6 ms, and bandwidth 290 Hz/pixel.

#### MEG Preprocessing

Raw MEG recordings were first processed with Max-Filter (version 2.2.15). Signal-space separation and temporal signal-space separation were used to suppress external interference, with continuous movement compensation based on the head-position-indicator coils. During this step, data were downsampled from 1000 to 250 Hz and a correlation limit of 0.98 was used to separate the internal and external signal subspaces.

The data were converted to Statistical Parametric Mapping format and processed in MATLAB using the Oxford Centre for Human Brain Activity Software Library and custom LBPD routines, incorporating functionality from SPM, FieldTrip, and FSL. Continuous recordings were visually inspected for large artefacts. Independent component analysis was then used to identify ocular and cardiac components by their relation to the EOG and ECG channels and by their characteristic spatial patterns; confirmed artefactual components were removed before signal reconstruction.

The cleaned data were segmented from 100 ms before to 3400 ms after excerpt onset and baseline-corrected relative to the 100-ms prestimulus interval. Trials containing residual artefacts were marked as bad. Correct-response trials remaining after these quality exclusions were organized by melody length and as memorized, early-change, or late-change trials before trial averaging and source reconstruction.

#### Source Reconstruction

Source activity was reconstructed with a linearly constrained minimum-variance beamformer. Individual MEG data were co-registered to the corresponding T1-weighted MRI using the digitized head shape, anatomical landmarks, and coil positions. A single-shell forward model was constructed on an 8-mm grid in standard space; when an individual anatomical scan was unavailable, the MNI152 T1 template was used. Lead fields were reduced to a single orientation at each location, and normalized beamformer weights were applied independently at every time point. Sensor covariance was estimated from trials pooled across the experimental conditions. This yielded condition-specific, trial-averaged time series at 3559 brain locations from −0.1 to 3.4 s.

### BROAD-NESS Network Estimation

Broadband brain networks were estimated from the source-reconstructed time series using the PCA implementation of BROAD-NESS. The decomposition was performed independently for M3, M5, and M7 so that the network structure at each melody length was estimated from its own retained participant sample. Within each melody length, source activity was first averaged across participants and across the three conditions. Principal component analysis was then applied to the resulting matrix of 3559 source time series. The components were ordered by explained variance and their signs were oriented so that the source with the largest absolute loading had a positive weight.

The group-level component weights defined the spatial pattern of each network. Participant- and condition-specific activation time series were then reconstructed by projecting each participant’s original source activity onto these common weights. Thus, all participants and conditions within a melody length were represented in the same component space, while retaining their individual temporal dynamics.

To estimate the number of relevant brain networks captured by the decomposition, we computed the effective dimensionality (ED) of the eigenspectrum using the participation ratio:

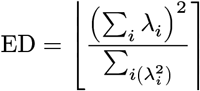

where *λ*_*i*_ is the variance explained by the *i*-th component. ED summarizes how many components contribute materially to the eigenspectrum without imposing an arbitrary cumulative-variance threshold.^18,19^ ED was two for M3 and three for both M5 and M7. Effective dimensionality was calculated directly from each decomposition at analysis time and determined the components used for phase-space reconstruction and matched RMS amplitude. The first three components were retained for the activation time-series overview and activation–behaviour control analysis at every melody length.

### Activation Time-Series Analysis

For each network and melody length, participant-level activation time series were compared between memorized and early-change trials and between memorized and late-change trials. At each time point, the within-participant condition difference was assessed with a two-sided paired-samples *t*-statistic. Temporally adjacent samples exceeding the two-sided cluster-forming threshold of *p* < .05 were grouped into clusters, and cluster mass was defined as the sum of the absolute *t*-statistics within the cluster.

Statistical significance was assessed with 2000 sign-flip permutations. On each permutation, the sign of the complete condition-difference time series was reversed for a random subset of participants, preserving its temporal dependence, and the largest cluster mass was retained. Observed clusters were compared with this maximum-statistic null distribution and considered significant at *p* < .05. As a complementary analysis, pointwise paired tests were corrected across time using FDR correction; these results are reported in the Supplementary Materials.

### Recurrence Quantification Analysis

The joint dynamics of the BROAD-NESS networks retained by effective dimensionality were characterized using phase-space reconstruction and recurrence quantification analysis (RQA). For each participant and condition, the activation time series of the retained components were treated as orthogonal axes of the state space. The reconstruction was therefore two-dimensional for M3 and three-dimensional for M5 and M7. Each sample represented the system’s network state at one time point. Analyses began at stimulus onset and extended until 800 ms after the final tone onset, giving windows of 0–1.5 s for M3, 0–2.2 s for M5, and 0–2.9 s for M7.

For every pair of time points, Euclidean distance was calculated between their phase-space coordinates. A pair was classified as recurrent when its distance was below 10% of the maximum distance observed for that participant and condition. Diagonal and vertical lines were required to contain at least two samples. Eight standard RQA measures were extracted: recurrence rate (RR), the proportion of recurrent points; mean diagonal line length (*L*), reflecting temporal predictability; determinism (DET), the proportion of recurrent points forming diagonal lines; diagonal-line entropy (ENTR), reflecting the diversity of diagonal durations; trapping time (TT), the mean vertical line length; laminarity (LAM), the proportion of recurrent points forming vertical lines; maximum vertical line length (*V*_max_), the longest period of state persistence; and divergence (DIV), the inverse of the longest diagonal line.

RQA metrics were calculated independently for each participant, melody length, and condition. Participants with RR equal to zero in any condition were excluded from all RQA analyses because their trajectories contained no recurrence points.

### RQA Statistical Analysis

Each RQA metric was analysed with a 3 × 3 repeated-measures ANOVA containing condition and melody length as within-participant factors. These models tested the main effects of condition and melody length and their interaction. FDR correction was applied across the eight RQA metrics separately for each omnibus effect. Partial *η*^2^ was used as the omnibus effect-size measure.

Follow-up analyses included paired comparisons among the three conditions, condition comparisons performed separately within each melody length, and the linear change from M3 to M7. Additional one-factor repeated-measures ANOVAs assessed melody-length effects within each condition. Paired tests used only participants with finite values for both observations and were summarized with Cohen’s *d*_*z*_. Cross-melody analyses used complete cases across all observations entering the relevant model.

A sensitivity analysis repeated the RQA calculations at 5%, 7.5%, 12.5%, and 15% of the participant-and condition-specific maximum distance, using the same exclusion rule and inferential models; the prespecified 10% analysis remained primary.

### RQA-Behaviour Associations

Associations between network dynamics and behavioural performance were assessed with two-sided Pearson correlations between each RQA metric and either response accuracy or mean reaction time across registered responses; no-response trials were excluded from the reaction-time means. Correlations were first calculated separately for each melody-length and condition combination. To characterize broader patterns, values were also averaged within participant across melody lengths for each condition and across conditions for each melody length before correlation. For each comparison and behavioural outcome, FDR correction was applied across the eight RQA metrics.

As an additional repeated-measures analysis, mixed models tested the overall RQA–behaviour association and whether this association varied by condition or melody length. Each behavioural outcome was modelled from condition, melody length, their interaction, the standardized RQA value, the RQA-by-condition interaction, and the RQA-by-melody-length interaction, with a random intercept for participant. Degrees of freedom were estimated using the Satterthwaite approximation. For each model term and behavioural outcome, FDR correction was applied across the eight RQA metrics.

### Activation-Behaviour Control Analysis

To test whether network activation amplitude adequately accounted for behavioural performance, activation was correlated across participants with accuracy and mean reaction time across registered responses at every time point, separately for each of the first three networks, melody length, and condition. All three networks were evaluated at every melody length so that the activation control used a common dimensional scope independent of the effective dimensionality selected for RQA. Pearson correlations were tested two-sided, and FDR correction was applied across all time points within each network-by-condition-by-outcome series.

For a formally matched test, activation magnitude was summarized as the root-mean-square (RMS) amplitude across the same retained components and time interval used for RQA in each participant, melody length, and condition. Accuracy was modelled from condition, melody length, their interaction, and standardized RMS activation amplitude, with a random intercept for participant. The primary inferential test compared this RMS-amplitude-adjusted model with a model additionally containing all eight standardized RQA measures, using maximum-likelihood estimation and an eight-degree-of-freedom likelihood-ratio test. Incremental marginal *R*^2^ quantified model fit. Metric-level coefficients and one-degree-of-freedom drop-one tests were treated as descriptive because of collinearity among the RQA measures, with FDR correction across the eight tests.

Predictive generalization was evaluated with nested participant-wise ridge regression. The outer cross-validation left one participant out, and three-fold participant-grouped cross-validation within each outer training set selected the ridge penalty from 29 logarithmically spaced values between 10^2^ and 10^−5^. Condition, melody length, their interaction, and RMS activation amplitude were unpenalized; only the eight RQA measures were regularized. Continuous predictors were standardized using parameters estimated within the relevant training fold. Prediction uncertainty was quantified by 10,000 participant-level bootstrap resamples of the paired outer-fold errors, and a two-sided paired sign-flip test used 100,000 permutations.

### Software and Reproducibility

MEG preprocessing and source reconstruction were performed in MATLAB R2016b using MaxFilter 2.2.15, OSL 2017Sep06 (`osl-core` commit `cf79690`), SPM12 revision 6906, FSL 5.0.9, the bundled FieldTrip distribution, and custom LBPD routines. All analyses following BROAD-NESS network estimation were performed in MATLAB R2026a Update 5 using BROAD-NESS and custom scripts; Connectome Workbench 2.2.1 was used for cortical visualization. The experiment was presented using PsychoPy. The code is available at the following link: https://github.com/DieSache/2026-rqa-auditory-recognition. The multimodal neuroimaging data will be provided upon reasonable request.

## Results

### Experimental and Analytical Overview

Participants judged whether three-, five-, and seven-tone excerpts were memorized or contained a single early or late tone change (Figure 1). We first characterized response accuracy, errors, omissions, and reaction times. Source-reconstructed MEG activity was then decomposed separately for each melody length with BROAD-NESS, yielding low-dimensional network activation time series shared across conditions and participants. We compared these activation time series between memorized and changed melodies and reconstructed joint trajectories from the two, three, and three networks retained for M3, M5, and M7, respectively. Recurrence quantification analysis (RQA) was used to test whether the temporal organization of these trajectories differed between conditions and whether individual differences in recurrent dynamics were related to behavioural performance.

### Behavioural Performance Varied with Change Position

Correct-response proportions differed among memorized, early-change, and late-change melodies (*F* (2, 104) = 8.8, FDR-adjusted *p* < .001, partial *η*^2^ = 0.145), varied with melody length (*F* (2, 104) = 4.03, FDR-adjusted *p* = 0.02, partial *η*^2^ = 0.072), and showed a condition-by-melody-length interaction (*F* (4, 208) = 9.04, FDR-adjusted *p* < .001, partial *η*^2^ = 0.148; Figure 2). Aggregate response proportions are shown in Figure S1. For M3 and M5, early changes produced more correct responses than both memorized and late-change melodies. For M7, late changes produced more correct responses than both memorized and early-change melodies. The complementary analysis of incorrect responses showed a condition effect (*F* (2, 104) = 14.83, FDR-adjusted *p* < .001, partial *η*^2^ = 0.222) and a condition-by-melody-length interaction (*F* (4, 208) = 7.9, FDR-adjusted *p* < .001, partial *η*^2^ = 0.132). No-response proportions were comparatively stable: neither condition, melody length, nor their interaction was significant (all FDR-adjusted *p* ≥ 0.08).

**Figure 2:**
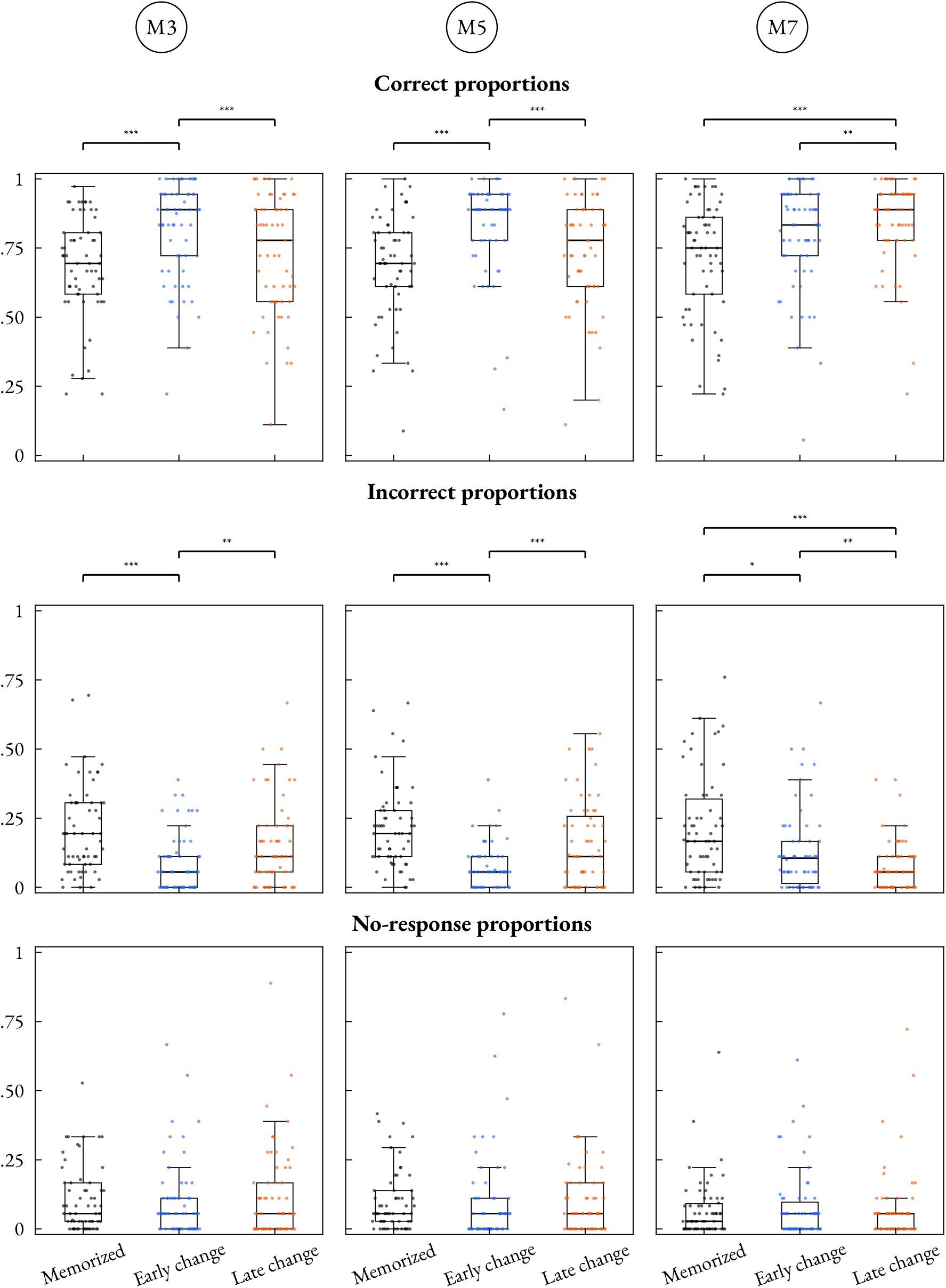
Behavioural response proportions across melody length and condition. Columns show M3, M5, and M7 melodies, and rows correct, incorrect, and no-response proportions. Boxplots and subject-level points show distributions for 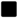 memorized (M), 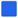 early-change, and 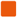 late-change trials. Brackets mark FDR-corrected condition comparisons. ∗ *p* < .05, ∗ ∗ *p* < .01, ∗ ∗

Reaction times for correct responses showed effects of condition (*F* (2, 104) = 38.13, FDR-adjusted *p* < .001, partial *η*^2^ = 0.423), melody length (*F* (2, 104) = 2172.9, FDR-adjusted *p* < .001, partial *η*^2^= 0.977), and their interaction (*F* (4, 208) = 5.37, FDR-adjusted *p* < .001, partial *η*^2^ = 0.094; Figure S2). The large melody-length effect reflected the later completion of longer excerpts when reaction time was measured from excerpt onset. Within melody lengths, correct responses to early changes were faster than responses to late changes for M3, M5, and M7; early changes were also faster than memorized melodies for M5 and M7. Only 17 participants had finite incorrect-response reaction-time means in all nine condition-by-melody-length combinations; the incorrect-response RT ANOVA was therefore based on these 17 complete cases. Incorrect-response times increased with melody length (*F* (2, 32) = 158.21, FDR-adjusted *p* < .001, partial *η*^2^ = 0.908), but showed no condition effect or interaction (both FDR-adjusted *p* = 0.4). Correct responses were faster than incorrect responses in 7 of the nine melody-length and condition combinations after correction for the displayed comparisons. The underlying behavioural descriptive statistics are reported in Table 2.

**Table 2:** Behavioural descriptive statistics. Response values are participant-level mean percentages ± SD (*n* = 53). Reaction-time values are mean milliseconds ± SD, with the number of participants contributing a finite mean in parentheses.

| Melody | Condition | Correct (%) | Incorrect (%) | No response (%) | Correct RT (ms) | Incorrect RT (ms) |
| --- | --- | --- | --- | --- | --- | --- |
| M3 | Memorized | 68.3 ± 17.9 | 20.2 ± 16.3 | 11.5 ± 12 | 1654 ± 197 (61) | 1917 ± 354 (58) |
|  | Early change | 80.2 ± 18 | 9.3 ± 10.7 | 10.5 ± 14.4 | 1592 ± 271 (61) | 1830 ± 462 (37) |
|  | Late change | 71.3 ± 21.8 | 16.1 ± 15.6 | 12.6 ± 17.1 | 1700 ± 238 (61) | 2009 ± 392 (48) |
| M5 | Memorized | 69.5 ± 17.2 | 20.2 ± 13 | 10.2 ± 10.9 | 2411 ± 246 (60) | 2679 ± 368 (58) |
|  | Early change | 82.6 ± 17 | 7.9 ± 7.8 | 9.5 ± 16.1 | 2227 ± 227 (60) | 2725 ± 431 (44) |
|  | Late change | 72.8 ± 20.4 | 16.6 ± 15.8 | 10.6 ± 16 | 2465 ± 228 (60) | 2556 ± 385 (44) |
| M7 | Memorized | 71.2 ± 19.7 | 21.5 ± 18.3 | 7.3 ± 11.1 | 3059 ± 236 (62) | 3269 ± 402 (57) |
|  | Early change | 76.8 ± 19.3 | 14.4 ± 15.5 | 8.8 ± 13.3 | 2902 ± 274 (62) | 3275 ± 381 (46) |
|  | Late change | 84.1 ± 15.9 | 8.1 ± 8.7 | 7.8 ± 13.9 | 3055 ± 194 (62) | 3232 ± 538 (42) |

### Melody-Specific Low-Dimensional Network Dynamics

The first three BROAD-NESS components explained 61.1%, 16.6%, and 6.9% of the variance in M3; 59.9%, 17.6%, and 6.5% in M5; and 54.4%, 20.1%, and 8.6% in M7. The corresponding effective dimensionality was 2 networks for M3, 3 for M5, and 3 for M7. Thus, despite differences in excerpt duration, the dominant source dynamics at each melody length were captured by a small number of distributed components.

Although BROAD-NESS was performed independently for each melody length, the resulting components showed qualitatively similar spatial patterns. The first component in each decomposition, which explained most of the variance, was dominated by bilateral primary and secondary auditory cortices together with medial and posterior cingulate regions. The second components retained bilateral auditory contributions but extended more strongly into memory-and prediction-related regions, including the hippocampal and inferior temporal cortices, insula, anterior cingulate cortex, and ventromedial prefrontal cortex. The third components explained substantially less variance and showed more heterogeneous temporal and frontotemporal distributions; they contributed to the effective state spaces for M5 and M7 but not M3. Across melody lengths, the reconstructed trajectories therefore described the joint evolution of decomposition-specific components encompassing dominant auditory–cingulate activity and additional medial temporal, insular, and prefrontal systems rather than anatomically unspecified dimensions.

Memorized and changed melodies followed overlapping activation time courses but diverged during discrete periods (Figure 3). Cluster-corrected differences were present in the first two networks at all three melody lengths. For M3, these differences spanned 0.48 to 2.37 s; M5 showed several intervals between 0.83 and 2.17 s. M7 differences extended from 1.54 to 2.79 s across the retained networks. No significant clusters were detected in the third network for M3 or M5. Exact cluster-corrected intervals are reported in

**Figure 3:**
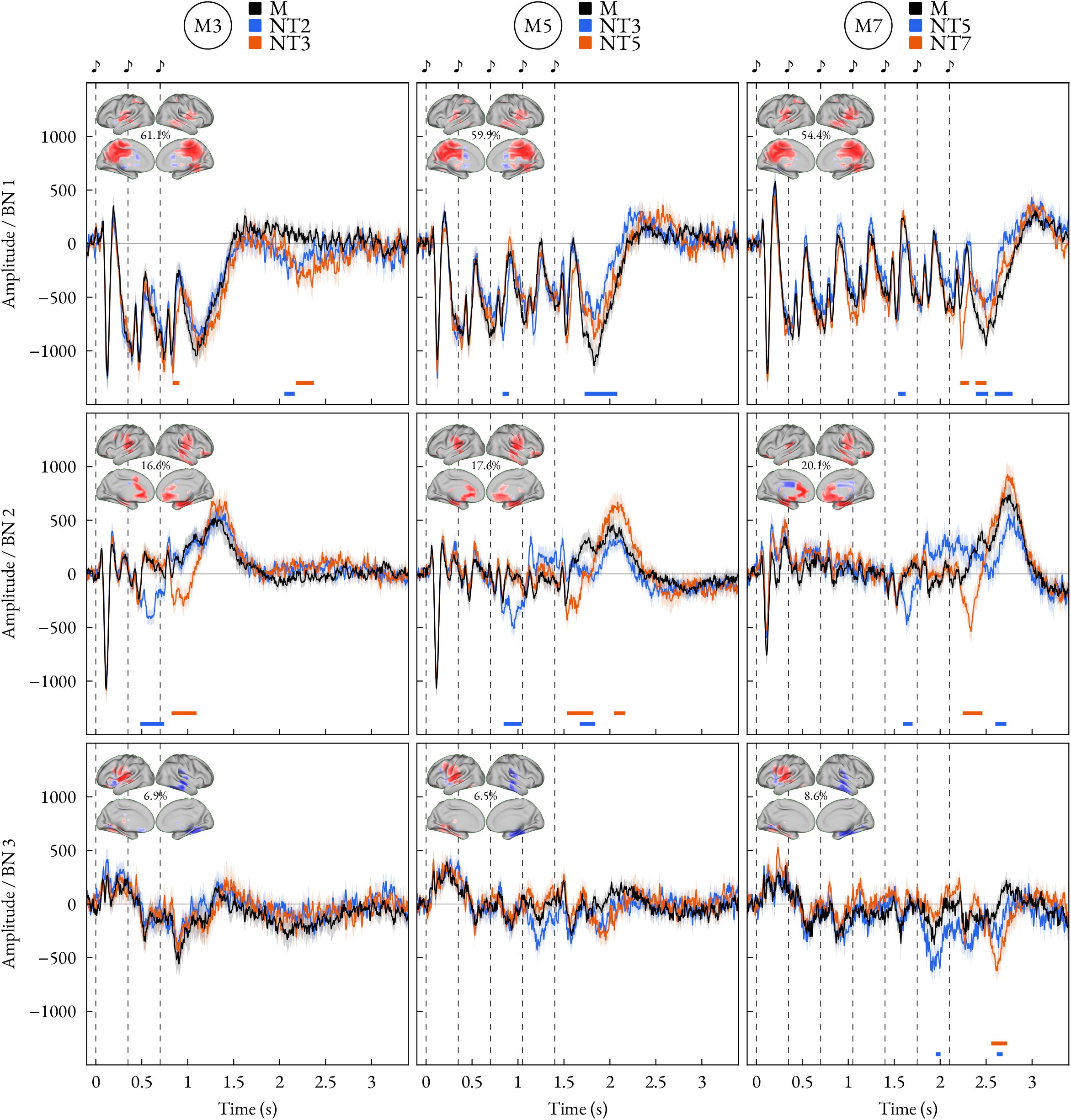
Cluster-corrected time series. Lines show mean activation time courses for 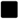 memorized (M), 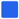 early-change, and 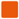 late-change melodies across brain-network clusters. Shaded bands show +/-standard error, dashed lines and musical notes bedded images show activation patterns with variance explained, and bottom bars mark significant early-and late-change windows.

Table S3. Pointwise FDR-corrected comparisons are shown in Figure S3.

### Memorized Melodies Followed More Structured Recurrent Trajectories

The geometry of the joint network trajectories differed consistently between memorized and changed melodies (Figure 4; descriptive and inferential statistics in Table 3). Condition affected all eight RQA metrics. Relative to both early- and late-change melodies, memorized melodies had a higher recurrence rate, longer diagonal and vertical structures, greater determinism, entropy, and laminarity, and lower divergence. The recurrence-rate effect was smaller than the effects for line structure (partial *η*^2^ = 0.083 versus 0.286−−0.443). Activity evoked by memorized melodies therefore returned more often to earlier states and, more prominently, formed longer, more persistent, and more predictable recurrence structures.

**Table 3:**
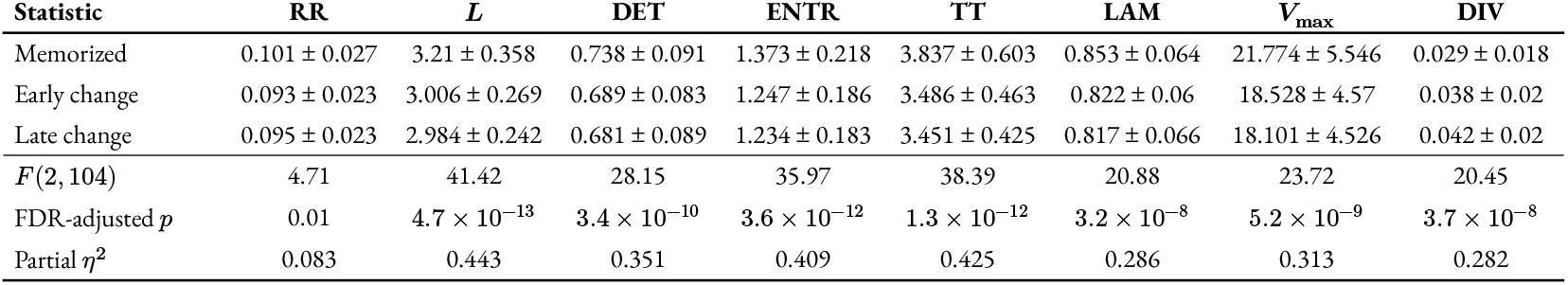
Recurrence quantification analysis (RQA) statistics by condition. RR, recurrence rate; *L*, mean diagonal line length; DET, determinism; ENTR, diagonal-line entropy; TT, trapping time; LAM, laminarity; *V*_max_, maximum vertical line length; DIV, divergence; SD, standard deviation; FDR, false-discovery rate. Descriptive values are participant-level means ± SD after averaging each metric across melody lengths for the 53 participants included in the cross-melody RQA analyses. Inferential rows report the main effect of condition.

**Figure 4:**
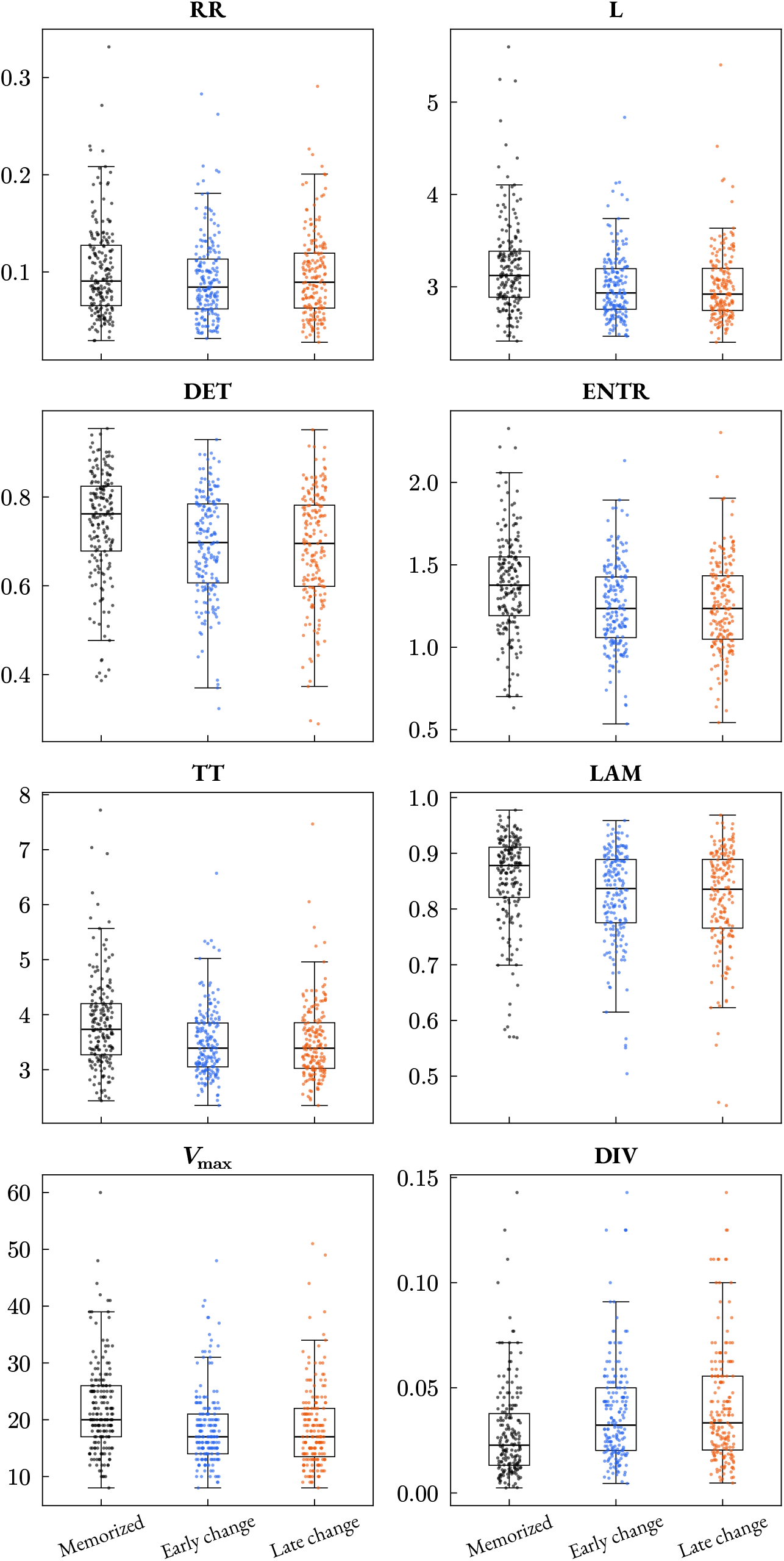
RQA metric distributions by condition family. Scatter and box plots show participant-level RQA metric distributions for 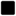 memorized (M), 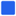 early-change, and 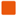 late-change melodies. Melody lengths are pooled within each condition family.

Condition effects were broadly stable across thresholds: six structural metrics remained significant from 5% to 15%, while *L* missed significance specifically at 5% and RR at 15% (full sensitivity results in Table S5).

All eight RQA metrics varied with melody length (all FDR-adjusted *p* ≤ 0.012). Recurrence rate and all line-structure measures were highest for M3, whereas divergence was lowest for M3 and highest for M5. These melody-length effects describe trajectories reconstructed using the effective dimensionality of each decomposition and therefore reflect both excerpt-specific dynamics and the corresponding two-, three-, and three-dimensional state spaces.

Condition-by-melody-length interactions survived correction for *L*, DET, ENTR, TT, and LAM (all FDR-adjusted *p* ≤ 0.012), but not for RR, *V*_max_, or DIV (all FDR-adjusted *p* ≥ 0.309). The memorized-versus-change distinction was present in the pooled condition contrasts for every metric, but its magnitude varied with melody length for several aspects of line structure. Condition-by-melody and melody-length distributions are provided in Figure S4 and Figure S10.

### Recurrent Organization Was Related to Behaviour

Participants with more structured recurrent trajectories tended to perform more accurately and respond more quickly (Figure 5). When values were averaged within melody length, accuracy correlated positively with the line-based RQA metrics across all three lengths (*r* = 0.329–0.504, all FDR-adjusted *p* ≤ 0.01) and negatively with divergence (*r* = −0.477 to −0.416, all FDR-adjusted *p* ≤ 0.001). The same pattern was present when values were averaged within condition: for memorized, early-change, and late-change melodies, accuracy increased with all line-based metrics (*r* = 0.399–0.55, all FDR-adjusted *p* ≤ 0.001) and decreased with divergence (*r* = −0.571 to −0.399, all FDR-adjusted *p* ≤ 0.001).

**Figure 5:**
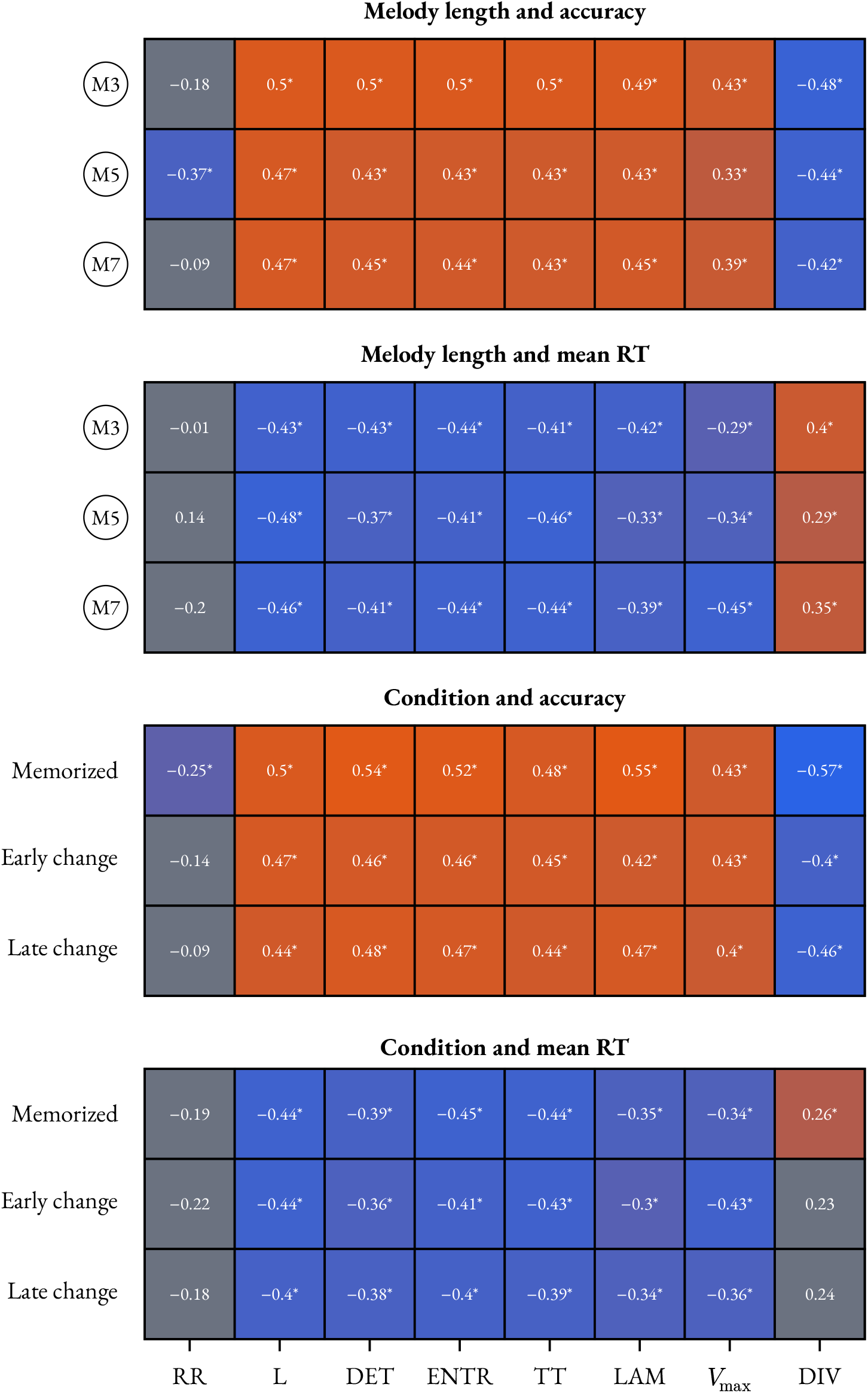
RQA-behaviour correlations across melody length and condition. Melody-length correlations were calculated after averaging each participant across conditions, whereas condition correlations were calculated after averaging across melody lengths. Heatmaps show Pearson correlations between RQA metrics and behavioural accuracy or mean response time across participants; no-response trials were excluded from mean response times. Tile values are Pearson correlation coefficients. Tile color indicates direction and magnitude, from 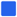 negative to 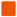 positive correlations; 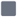 neutral tiles indicate non-significant correlations. Asterisks mark significant two-sided Pearson correlation tests, FDR-corrected across the eight RQA metrics.

Reaction-time associations had the complementary direction. Within melody lengths, greater line structure was related to faster responses (*r* = −0.476 to −0.286, all FDR-adjusted *p* ≤ 0.029), whereas greater divergence was related to slower responses (*r* = 0.286–0.403, all FDR-adjusted *p* ≤ 0.031). Across condition averages, all line-based associations survived correction for memorized, early-change, and late-change melodies (all FDR-adjusted *p* ≤ 0.022). Divergence was related to slower responses for memorized melodies (*r* = 0.257, FDR-adjusted *p* = 0.045), but narrowly missed correction for both change conditions. Recurrence rate showed no consistent relationship with either behavioural outcome. Condition-specific correlations are reported in Figure S8 and Figure S9.

In repeated-measures mixed models, the overall RQA slope survived FDR correction for accuracy for every RQA metric except RR. For reaction time, the overall slopes for *L* and TT also survived correction; no RQA interaction term survived correction. Because these models account for repeated observations but do not separate within-and between-participant RQA variation, the behavioural associations were clearest and most extensive for accuracy, while the reaction-time pattern was supported by both the aggregate correlations and the two corrected overall slopes.

### Recurrent Dynamics Explained Accuracy Beyond Matched RMS Activation Amplitude

Direct correlations between network activation amplitude and behaviour were sparse after correction across time (Figure S5, Figure S6, and Figure S7). The clearest effect was a negative association between first-network activation and reaction time during late-change M3 trials, which survived FDR correction at 81 time points; 16 samples in the same series were positively related to accuracy. Smaller M3 effects related early-change reaction time to BN1 and BN2 at 6 and 4 samples, respectively. For M5, 25 late-change samples in BN2 were related to reaction time. M7 effects were confined to memorized-trial reaction time in BN2 (1 sample), memorized-trial accuracy and reaction time in BN3 (14 and 14 samples), and early-change accuracy in BN3 (4 samples). No corrected time points were detected in the remaining network-by-condition series. Activation amplitude therefore showed localized associations but did not provide the broad account of behavioural performance observed for recurrent dynamics.

The formally matched analysis provided a direct model comparison. After adjustment for the condition-by-melody-length interaction and RMS activation amplitude, adding the eight RQA measures jointly increased marginal ***R***^2^ by 0.273 (*χ*^2^(8) = 154.49, *p* = 2.3 × 10^−29^). In nested participant-wise cross-validation, the ridge-regularized RQA model reduced held-out RMSE from 0.188 to 0.166, an improvement of 11.81% (participant-bootstrap 95% CI: 4.01%–17.65%; paired sign-flip *p* = 6 × 10^−4^). Prediction improved for 53 of 65 held-out participants. Thus, recurrent dynamics explained accuracy variation beyond whole-window RMS activation magnitude and improved prediction for held-out participants under regularization designed for the collinear RQA measures. The joint block test was the primary inference; descriptive metric-level mixed-model and ridge results are reported in Table S4.

## Discussion

The present study examined whether the recurrent organization of low-dimensional whole-brain activity relates to auditory recognition performance beyond activation magnitude. Memorized melodies elicited more structured recurrent trajectories, and participants with stronger recurrent organization were generally more accurate and faster. These dynamics explained variation in accuracy beyond the RMS activation amplitude of the same latent processes and improved prediction for held-out participants.

Behavioural performance depended on both melody length and the position of the altered tone. Early changes were recognized more accurately than memorized and late-change melodies in M3 and M5, whereas late changes were recognized most accurately in M7. Correct responses to early changes were faster than those to late changes at every melody length. This pattern places the neural results in the context of a temporally unfolding recognition decision: the evidence available to distinguish a changed sequence from a memorized one depends on when the alteration occurs and on the preceding sequence. The increase in reaction time with melody length also reflects the later completion of longer excerpts, because responses were timed from excerpt onset.

BROAD-NESS captured the dominant source dynamics with two latent networks for M3 and three for M5 and M7. Although the decompositions were estimated independently, their components showed qualitatively similar spatial patterns: dominant bilateral auditory and cingulate contributions were accompanied by medial temporal, insular, and prefrontal contributions in additional components. These patterns situate the trajectories within distributed systems relevant to auditory processing, memory, salience, and prediction, consistent with our earlier BROAD-NESS findings^38^. Memorized and changed melodies followed overlapping activation time courses but differed during discrete periods in the first two networks at each melody length. Thus, the decomposition revealed both a compact organization of distributed activity and sensitivity of its activation to the recognition conditions; RQA then characterized how the simultaneous activity of these networks was organized through time.

The RQA findings distinguished the frequency of recurrence from the temporal structure of recurrent trajectories. Memorized melodies produced a modest increase in recurrence rate, showing that their neural trajectories returned more often to previously occupied regions of state space. Recurrence rate, however, showed no consistent corrected association with behaviour. Performance was instead related to the structure of those returns. Participants who performed better exhibited longer and more deterministic diagonal patterns, greater vertical persistence and laminarity, and lower divergence. Diagonal-line entropy also increased alongside determinism and line length, suggesting that effective trajectories were neither random nor confined to one rigid sequence, but contained a varied repertoire of temporally ordered returns. Thus, the behaviourally relevant property was not simply how often the system revisited a state, but whether those returns formed sustained and coherent temporal structures.

The condition effects and individual differences address complementary aspects of recognition. Memorized melodies elicited more structured recurrence, but were not uniformly recognized more accurately than changed melodies. The neural condition contrast therefore cannot be read as a simple ranking of task difficulty. Rather, recurrence distinguished responses to unchanged and altered sequences, while the behavioural associations showed that participants with more structured trajectories tended to perform better across the recognition task.

This result does not imply that activation magnitude is unimportant. Evoked-response amplitude, oscillatory power, and response timing have repeatedly been associated with auditory memory and recognition^5,9,10,42–46^, and the present activation time series contained some localized behavioural associations. The more specific conclusion is that activation magnitude did not provide a sufficient account of the present data. In the formally matched comparison, adding the eight RQA measures to a model containing condition, melody length, their interaction, and RMS activation amplitude increased marginal variance explained by 0.273. The regularized model also reduced held-out prediction error by 11.81%. Because the RQA measures were strongly collinear, this evidence applies to their joint contribution rather than to any single metric considered independently. Nevertheless, both the model comparison and participant-wise cross-validation show that recurrent organization contained information about accuracy that was not captured by matched whole-window RMS amplitude.

Our targeted literature screen places this contribution in the context of how neural activity has been related to behaviour. Among the 59 retained electrophysiological studies of memory, recognition, and sequence learning, 71% used evoked-response magnitude or timing or oscillatory magnitude (Table S1). Other studies established behavioural relationships with functional connectivity, neural entrainment, intersubject synchronization, and multivariate representations^47–54^, while our earlier BROADNESS study demonstrated the behavioural relevance of temporal-state dynamics^38^. Together with research characterizing low-dimensional neural organization, this literature motivates a direct test of whether the joint dynamics of latent processes explain performance beyond their activation strength. The present matched comparison addresses that question: recurrent organization accounted for individual differences in recognition accuracy beyond the RMS amplitude of the same latent processes. This connects the description of low-dimensional brain dynamics to an explanation of behavioural variation and identifies temporal organization as a source of information that activation magnitude alone does not capture.

These findings are consistent with the two linked principles of the low-dimensional hypothesis for cognition^15^. The source-reconstructed MEG activity was dominated by a small number of latent networks, while individual behaviour was associated with the organization of their joint trajectory. The second observation is critical because low dimensionality alone does not explain cognition: a compact set of processes may be present across participants and conditions while differing substantially in how it evolves. Low dimensionality and dynamical organization therefore describe complementary aspects of neural activity: the former constrains the set of processes through which distributed activity is expressed, whereas the latter captures how those processes combine and reorganize through time^24,25^. Within the integrated BROAD-NESS framework^38^, this distinction can be examined directly because the latent networks define the principal processes organizing whole-brain activity while their simultaneous evolution defines the trajectory of the system. The present findings suggest that the cognitive relevance of low-dimensional organization lies not only in reducing neural activity to a compact set of processes, but critically in the repertoire of temporal relationships that can emerge among them. In this view, cognitive performance might not simply depend on which latent processes are recruited or how strongly they are activated, but on how their relationships are dynamically organized as cognition unfolds.

Melody recognition is particularly informative in this respect because every tone is interpreted in the context established by the preceding sequence^34,55–57^. An unchanged continuation can remain compatible with a learned temporal structure, whereas an altered tone changes that context and may redirect the system into a different region of its joint state space. Consistent with this interpretation, a single changed tone was associated with shorter and less deterministic recurrence structures, reduced persistence, and greater divergence. MEG makes this rapidly unfolding organization accessible at the timescale of the sequence. Because RQA summarized an extended analysis window, however, the present results do not localize the recurrent reorganization to the precise moment of the altered tone. The prediction-error interpretation is therefore plausible and consistent with previous findings in auditory predictive processing^34,38^, but it remains to be tested directly with time-resolved or event-specific trajectory analyses.

The stimuli were drawn from a single Western classical musical piece. Whether the same relationships between recurrent organization and recognition performance hold for other musical traditions remains to be established.

Several limitations should be acknowledged to contextualize the current results and define directions for future work. Neural trajectories were derived from condition averages of correct-response trials, and the behavioural associations therefore capture between-participant relationships between performance and the dynamics expressed during successful recognition, rather than trial-level predictors of individual responses. Extending the framework to singletrial trajectories will be important for determining whether recurrent organization predicts recognition success from trial to trial and how such organization develops around behaviourally relevant events. Similarly, the observational design establishes robust associations between joint neural dynamics and behaviour but does not determine whether these dynamics play a causal role in performance, an important question for future experimental and perturbational studies. Strong collinearity among RQA measures was very much expected and does not represent a problem in itself, but further means that their behavioural contribution is most appropriately interpreted as evidence for recurrent organization as a whole rather than for unique effects of individual metrics. Nested participant-wise cross-validation demonstrated that this organization improved prediction in held-out participants within the present cohort, providing evidence that the behavioural relationship extends beyond in-sample model fit; however, validation in independent cohorts is called to provide the next, even stronger test of its generalizability. Finally, the sample included Danish and Chinese participants living in Denmark, offering an opportunity for future work to examine whether cultural or experiential background shapes low-dimensional neural dynamics and their relationship to cognition. Despite its relevance often being overlooked, we advocate for the importance of replication across independent cohorts, stimulus sets, and cognitive domains, together with trial-resolved and temporally localized analyses, to establish how broadly the present strong principles generalize.

In conclusion, melody recognition was associated with the recurrent temporal organization of lowdimensional whole-brain activity. More structured dynamics accompanied better performance and explained behavioural variation beyond matched activation magnitude. These findings suggest that the cognitive significance of low-dimensional neural organization lies not only in which processes are engaged or how strongly, but in how their relationships evolve over time. Understanding cognition may therefore require explaining both the processes that organize distributed neural activity and the dynamics that emerge from their joint evolution.

## Acknowledgment

The Center for Music in the Brain (MIB) is funded by the Danish National Research Foundation (project number DNRF117), The Lundbeck Foundation (R469-2024-1573) and Købmand Herman Sallings Fond.

L.B. is supported by Sapere Aude: Independent Research Fund Denmark (DFF) Research Leader (grant ID: 10.46540/5253-00003B), Lundbeck Foundation (Talent Prize 2022), Carlsberg Foundation (CF20-0239), Center for Music in the Brain, Linacre College of the University of Oxford and Nordic Mensa Fund.

MLK is supported by Center for Music in the Brain and Centre for Eudaimonia and Human Flourishing, which is funded by the Pettit and Carlsberg Foundations.

## Author contributions

LB, KFC and MR conceived the hypotheses. LB and GFR designed the study and created the stimuli. LB, MLK and PV recruited the resources for the experiment. MK, KFC, SS and MS collected the data. KFC, LB and MK performed pre-processing, computed BROAD-NESS, RQA and the statistical analysis. MLK, GFR, MR and PV provided essential help to interpret and frame the results within the neurosci-entific literature. KFC and LB wrote the first draft of the manuscript. KFC, GFR and LB prepared the figures. All the authors contributed to and approved the final version of the manuscript.

## Competing interests’ statement

The authors declare no competing interests.

## Supplementary materials

### Neural–behaviour literature screen

We conducted two complementary, method-neutral PubMed searches for human electrophysiological studies that related neural measurements directly to memory, recognition, or learning behaviour. Both searches were performed on 6 August 2026 and restricted to records published from 2015 onward. The auditory search returned 143 records. The complementary search for sequential, continuous, or otherwise temporally unfolding stimuli returned 67 records. 2 records occurred in both searches, yielding 208 unique records for title-and-abstract screening.

A study was retained only when it explicitly tested a statistical relationship between a neural measure and observed behavioural performance. Eligible relationships included comparisons between correct and incorrect or subsequently remembered and forgotten trials, participant-level neural–behaviour correlations, and models predicting accuracy, response time, recall, discrimination, or learning; explicit null tests were retained. We excluded studies that reported neural and behavioural condition effects without testing their relationship, decoded content without relating the neural measure to performance, or related neural activity only to an external cognitive, clinical, or demographic measure. Non-human studies and studies without a relevant behavioural outcome were also excluded. Retained measures were classified as evoked-response magnitude or timing, oscillatory magnitude, neural coordination, multivariate representation or prediction, or temporal-state dynamics. Neural coordination comprised phase, entrainment, connectivity, synchronization, and cross-frequency coupling; multivariate methods comprised decoding, representational similarity, and predictive models using distributed neural features. For studies with several analyses, classification followed the measure tested against behaviour.

59 studies met the direct neural–behaviour criterion: 45 used auditory or musical tasks and 14 used other temporally unfolding tasks. Evoked-response magnitude or timing was used in 20 studies and oscillatory magnitude in 22 studies. Together, these accounted for 42 of 59 retained studies (71%). 8 studies used neural-coordination measures, 8 used multivariate representations or prediction, and 1 used temporal-state dynamics. No retained study used a standalone entropy or information-complexity measure as its primary behavioural neural variable. These findings characterize the retained PubMed sample and are not intended as an exhaustive systematic review.

**Table S1:**
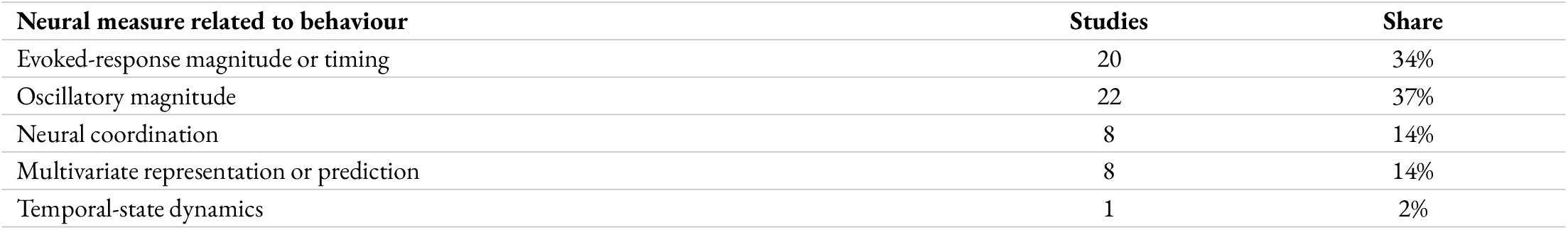
Methodological distribution of studies directly relating neural measurements to task performance. Percentages use all 59 retained studies as the denominator.

**Exact auditory query**.

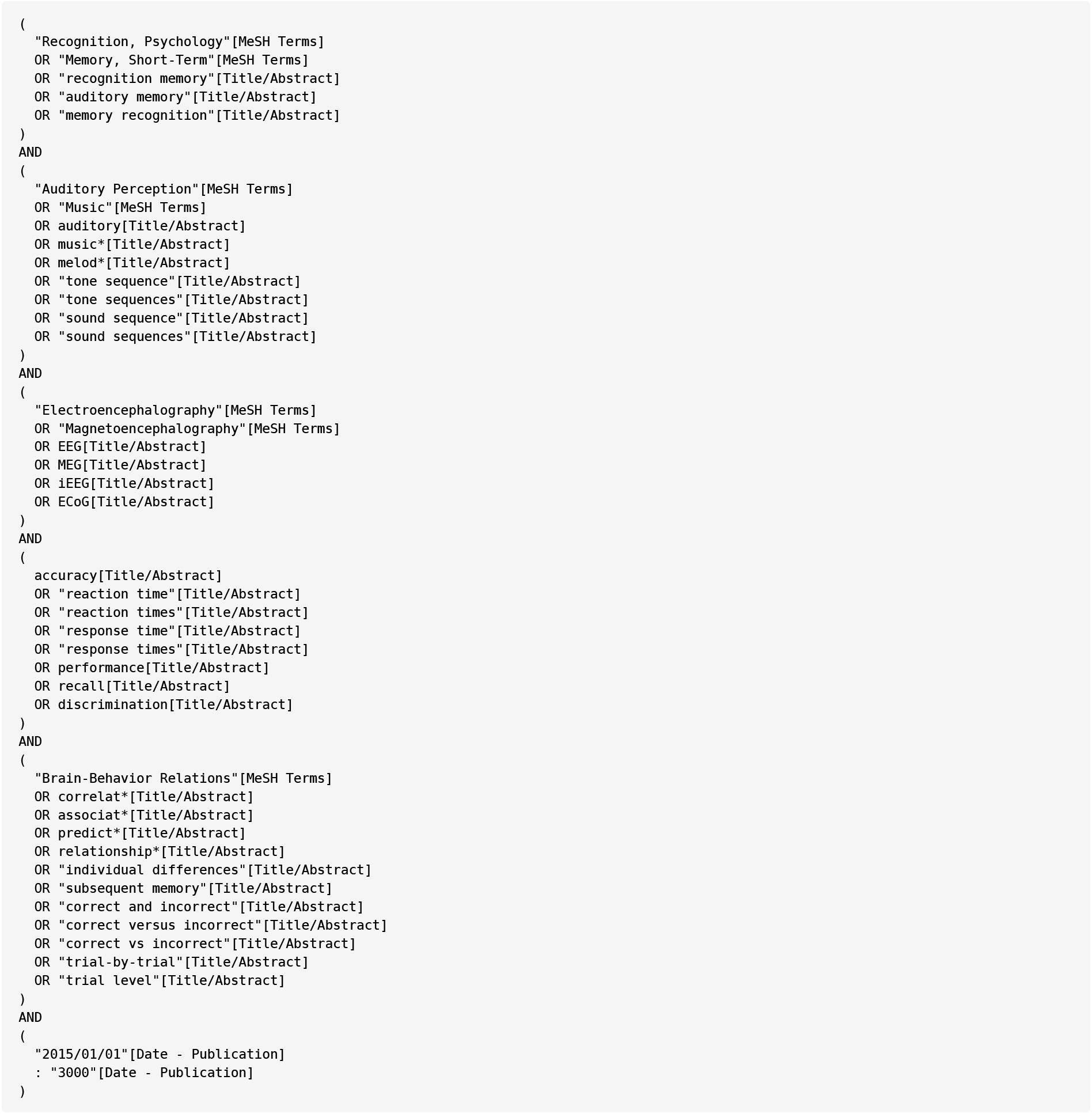

**Exact temporal-stimulus query**.

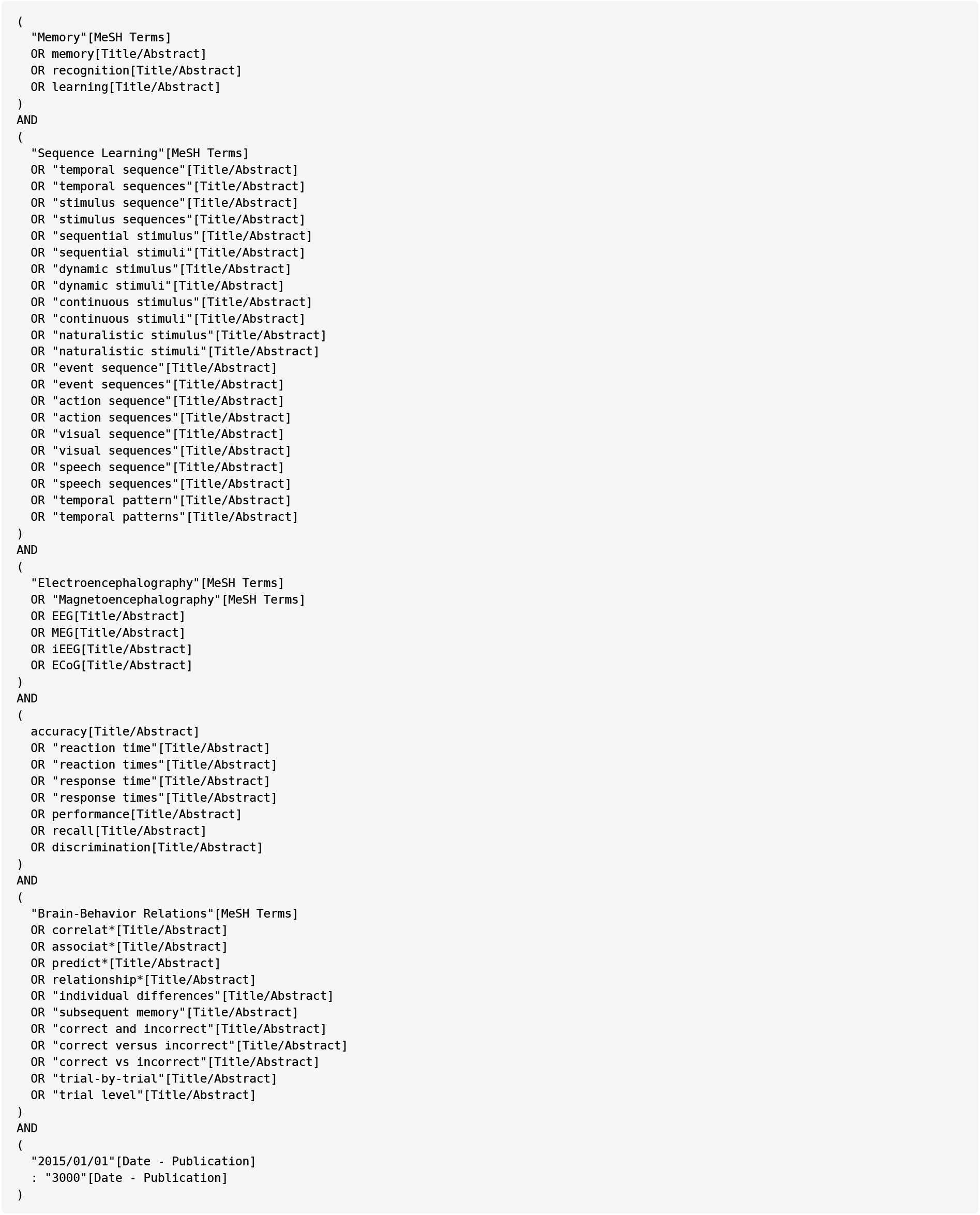

**Table S2:** Direct neural–behaviour evidence map. PMID, PubMed identifier.

| Study | PMID | Task domain | Neural measure | Behavioural measure |
| --- | --- | --- | --- | --- |
| **Recent auditory and musical studies (2020–2026)** |  |  |  |  |
| Jiang et al. (2026) | 42224926 | Auditory working memory | Evoked-response magnitude/<br>timing | Accuracy and response time<br>(ratio) |
| Lin et al. (2026) | 41934791 | Lexical-tone recognition in noise | Evoked-response magnitude/<br>timing | Accuracy |
| Beckers et al. (2025) | 39719815 | Auditory working memory | Oscillatory magnitude | Accuracy |
| Bonetti et al. (2025) | 41020476 | Auditory recognition memory | Temporal-state dynamics | Accuracy and response time |
| Chow et al. (2025) | 39689526 | Auditory recognition memory | Evoked-response magnitude/<br>timing | Accuracy (d-prime) |
| Cui et al. (2025) | 41176206 | Auditory working memory | Evoked-response magnitude/<br>timing | Response time |
| Del Popolo Cristaldi et al.<br>(2025) | 40136269 | Visual recognition memory | Oscillatory magnitude | Accuracy (level and dual-task<br>difference) |
| Hebron et al. (2025) | 40928755 | Visual working memory | Oscillatory magnitude | Accuracy (d-prime difference) |
| Pashkov et al. (2025) | 40317131 | Auditory working memory | Neural coordination | Recall score (normalized digits) |
| Prince et al. (2025) | 40561110 | Auditory working memory | Oscillatory magnitude | Response time |
| Tian et al. (2025) | 39215886 | Auditory working memory | Oscillatory magnitude | Accuracy and response time |
| Zhao et al. (2025) | 41213442 | Audiovisual working memory | Evoked-response magnitude/<br>timing | Accuracy |
| Borderie et al. (2024) | 38442128 | Auditory sequence working<br>memory | Neural coordination | Accuracy and response time<br>(inverse efficiency) |
| Frei et al. (2024) | 39444375 | Speech-in-noise recognition | Neural coordination | Accuracy |
| Inguscio et al. (2024) | 38608331 | Auditory verbal working<br>memory | Oscillatory magnitude | Accuracy |
| Kausel et al. (2024) | 38702896 | Audiovisual working memory | Oscillatory magnitude | Accuracy |
| Rao et al. (2024) | 39423321 | Visual working memory | Multivariate representation/<br>prediction | Recall and response time (pre-<br>post differences) |
| Cho et al. (2023) | 36653897 | Visual recognition memory | Multivariate representation/<br>prediction | Accuracy and response time |
| Paraskevoudi et al. (2023) | 35918912 | Auditory recognition memory | Evoked-response magnitude/<br>timing | Accuracy |
| Pomper et al. (2023) | 37691137 | Auditory short-term memory | Evoked-response magnitude/<br>timing | Accuracy |
| Qin et al. (2023) | 36866516 | Visual working memory | Oscillatory magnitude | Accuracy |
| Albouy et al. (2022) | 35196074 | Auditory working memory | Neural coordination | Accuracy (d-prime) |
| Bidelman et al. (2021) | 33323838 | Auditory working memory | Evoked-response magnitude/<br>timing | Accuracy (K capacity estimate) |
| Inguscio et al. (2021) | 34966418 | Verbal working memory | Oscillatory magnitude | Response time |
| Herman et al. (2020) | 32483253 | Auditory working memory | Oscillatory magnitude | Accuracy |
| Müller et al. (2020) | 29514397 | Speech-in-noise recognition | Neural coordination | Accuracy |
| **Earlier auditory and musical studies (2015–2019)** |  |  |  |  |
| Bastarrika-Iriarte et al. (2019) | 31015026 | Auditory recognition memory | Oscillatory magnitude | Accuracy (remembered/<br>forgotten) |
| Dehais et al. (2019) | 30508609 | Auditory alarm detection | Evoked-response magnitude/<br>timing | Accuracy |
| Morris et al. (2019) | 30553033 | Speech-in-noise perception | Evoked-response magnitude/<br>timing | Accuracy |
| Tune et al. (2018) | 29430736 | Selective auditory attention | Oscillatory magnitude | Accuracy |
| Wessel et al. (2018) | 27909006 | Visual working memory | Evoked-response magnitude/<br>timing | Accuracy |
| Zaske et al. (2018) | 28802769 | Auditory recognition memory | Oscillatory magnitude | Accuracy |
| Albouy et al. (2017) | 28343866 | Auditory working memory | Neural coordination | Accuracy (d-prime) |
| Impey et al. (2017) | 28864916 | Visual working memory | Evoked-response magnitude/<br>timing | Response time |
| Meltzer et al. (2017) | 28526621 | Sentence working memory | Oscillatory magnitude | Recall accuracy (cued) |
| Ou et al. (2017) | 28144832 | Lexical-tone perception and<br>production | Evoked-response magnitude/<br>timing | Production distinctiveness<br>(acoustic) |
| Yu et al. (2017) | 28720790 | Auditory duration working<br>memory | Oscillatory magnitude | Accuracy |
| Lefebvre et al. (2016) | 26903419 | Auditory sequence working<br>memory | Evoked-response magnitude/<br>timing | Accuracy (capacity estimate) |
| Mathias et al. (2016) | 27027544 | Musical recognition memory | Evoked-response magnitude/<br>timing | Accuracy |
| Sun et al. (2016) | 27973531 | Visual recognition memory | Multivariate representation/<br>prediction | Accuracy (remembered/<br>forgotten) |
| Toth et al. (2016) | 27421185 | Auditory figure-ground detection | Evoked-response magnitude/timing | Accuracy (d-prime) |
| Backer et al. (2015) | 25609643 | Auditory working memory | Oscillatory magnitude | Accuracy and response time (throughput) |
| Hawkins et al. (2015) | 25269110 | Phonological-form learning | Evoked-response magnitude/timing | Accuracy |
| Lim et al. (2015) | 26658862 | Auditory working memory | Oscillatory magnitude | Accuracy (psychometric-precision difference) |
| Wostmann et al. (2015) | 25390200 | Selective auditory attention | Evoked-response magnitude/timing | Accuracy |
| **Other temporally unfolding tasks** |  |  |  |  |
| He et al. (2026) | 41945608 | Visual sequence learning | Multivariate representation/prediction | Accuracy (sequence order) |
| Xue et al. (2026) | 41890133 | Associative episodic memory | Multivariate representation/prediction | Accuracy (d-prime) and recall rate (associative cued) |
| Dash et al. (2025) | 40938318 | Motor sequence learning | Multivariate representation/prediction | Typing speed (correct-sequence difference) |
| Mak et al. (2025) | 41161527 | Visual statistical learning | Oscillatory magnitude | Accuracy |
| Wu et al. (2022) | 36306242 | Visual episodic sequence memory | Multivariate representation/prediction | Recall accuracy (cued) |
| Lutz et al. (2021) | 33743012 | Visuomotor sequence learning | Oscillatory magnitude | Response time (random-sequence difference) |
| Zheng et al. (2021) | 33862254 | Visual sequence learning | Oscillatory magnitude | Response time (fixed-random difference) |
| Kucewicz et al. (2019) | 30847390 | Verbal episodic memory | Oscillatory magnitude | Recall status (recalled/forgotten) |
| Staudigl et al. (2019) | 31185803 | Cross-modal recognition memory | Multivariate representation/prediction | Accuracy |
| Crivelli-Decker et al. (2018) | 29753784 | Visual sequence learning | Oscillatory magnitude | Response time |
| Vila-Ballo et al. (2018) | 29339147 | Working-memory updating | Evoked-response magnitude/timing | Response time |
| Batterink et al. (2017) | 28324696 | Auditory statistical learning | Neural coordination | Response time (target detection) |
| Cohen et al. (2016) | 27844062 | Audiovisual narrative memory | Neural coordination | Accuracy |
| Donaldson et al. (2016) | 27059320 | Reversal learning | Evoked-response magnitude/timing | Error rate (post-reversal) |

**Figure S1:**
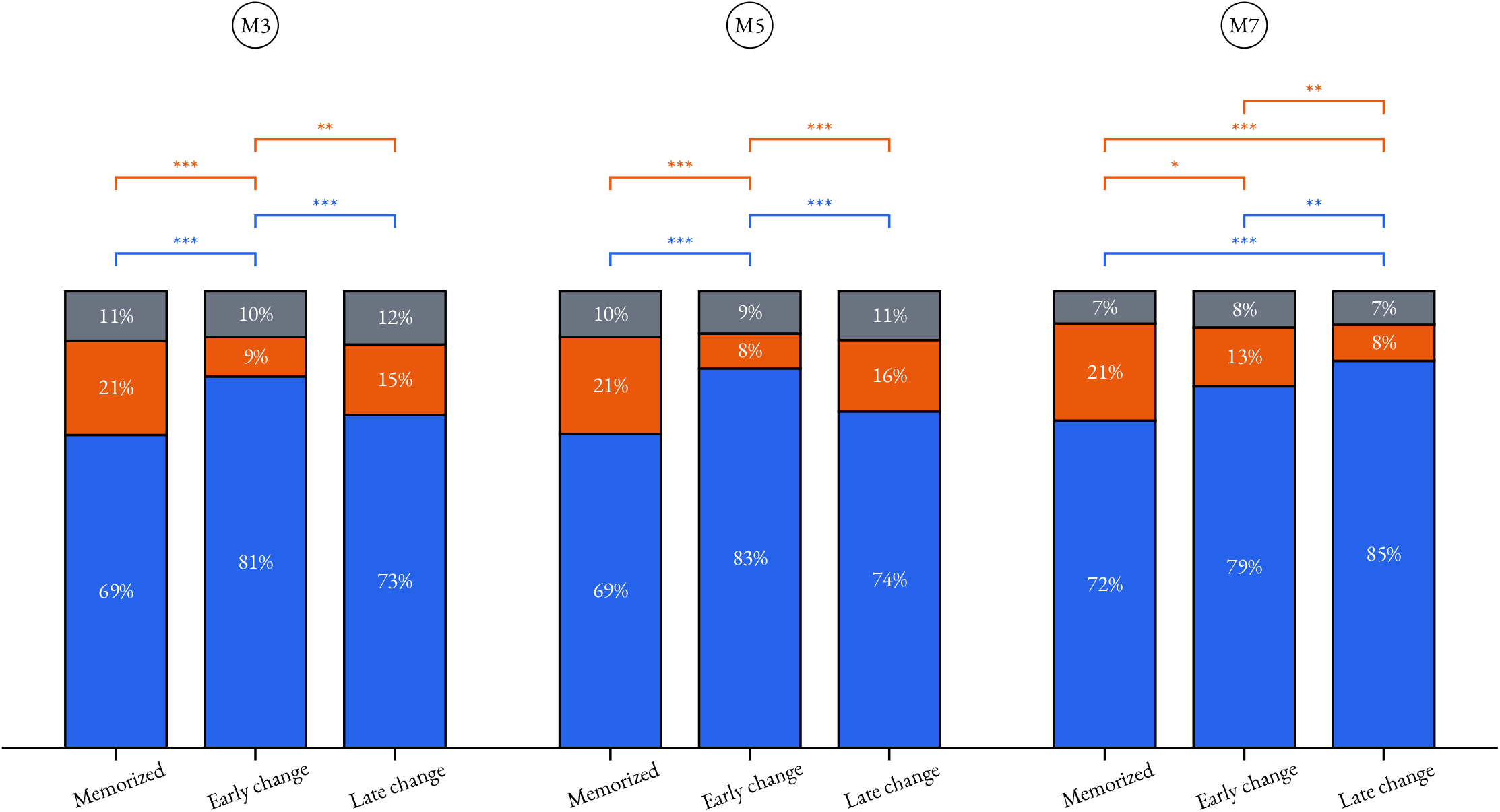
Behavioural response proportions across melody length and condition. Stacked bars show the proportion of 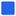 correct responses, 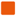 incorrect responses, and 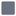 no responses across memorized (M), early-change, and late-change trials in the M3, M5, and M7 melody-length conditions. Percentages are shown within each response category. Brackets indicate significant FDR-corrected comparisons and use the color of the response category being compared. Asterisks denote significance levels: ∗ *p* < .05, ∗ ∗ *p* < .01, ∗ ∗ ∗ *p* < .001.

**Figure S2:**
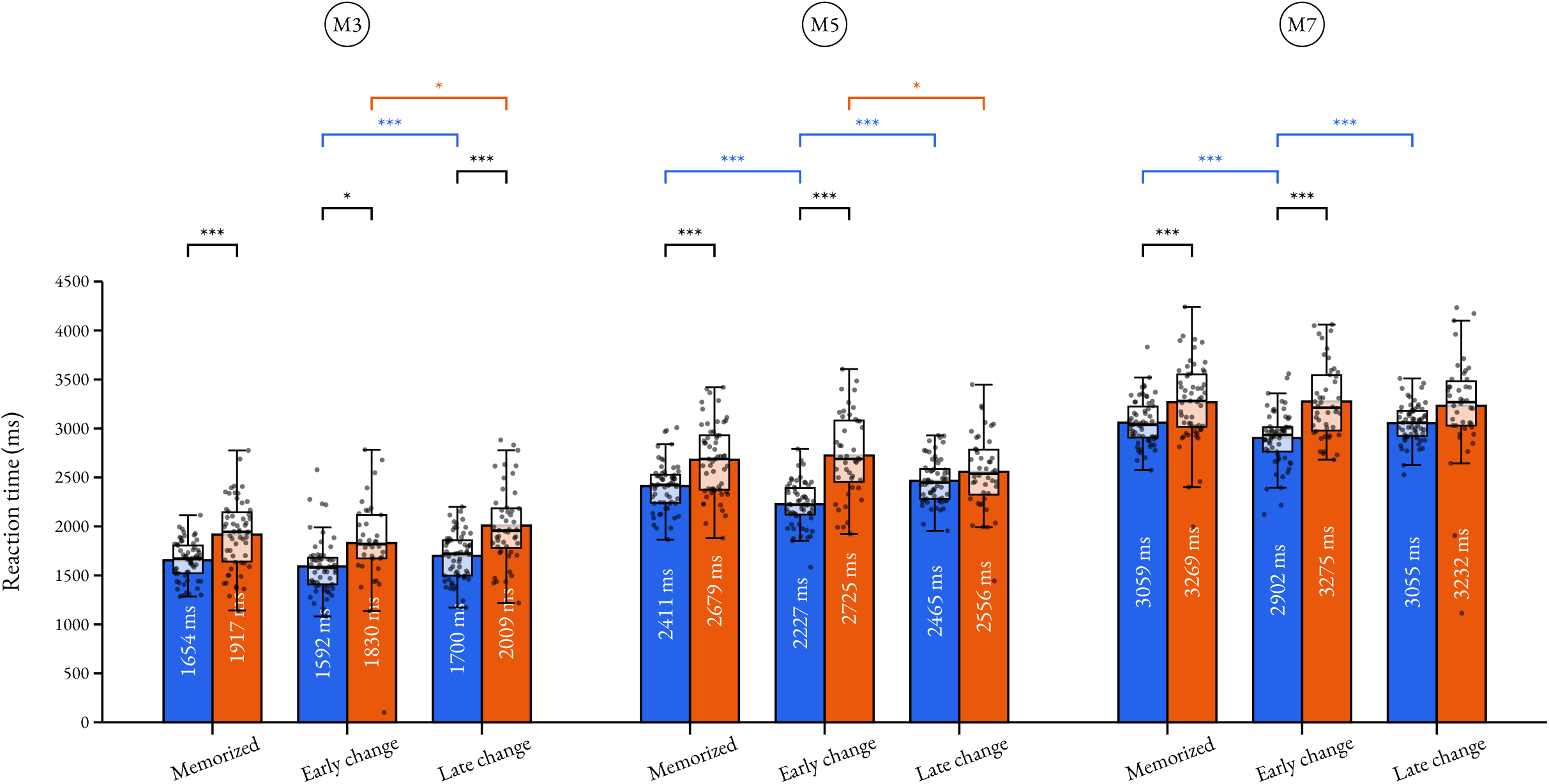
Reaction times across melody length and condition. Bars show mean reaction time in milliseconds for 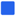 correct responses and 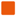 incorrect responses across memorized (M), early-change, and late-change trials in the M3, M5, and M7 melody-length conditions. Points show individual subject-level mean reaction times, and boxplots show medians, interquartile ranges, and 1.5-IQR whiskers. Black brackets compare correct with incorrect responses; brackets comparing conditions within one response category use that category’s color. Asterisks denote significance levels: ∗ *p* < .05, ∗ ∗ *p* < .01, ∗ ∗ ∗ *p* < .001.

**Figure S3:**
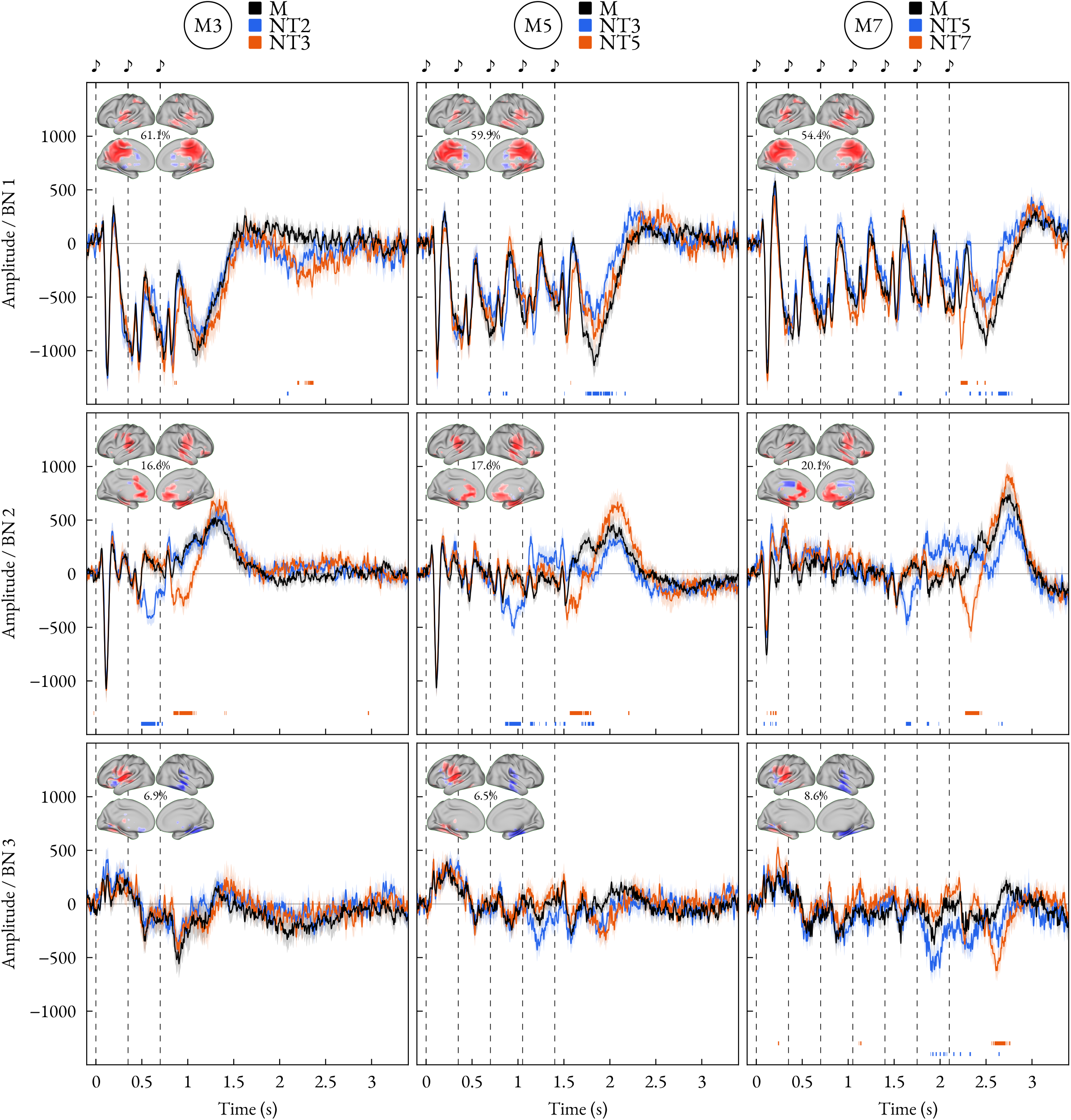
FDR-corrected time series. Lines show mean activation time courses for 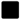 memorized (M), 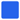 early-change, and 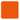 late-change melodies across brain-network clusters. Shaded bands show +/-standard error, dashed lines and musical notes mark tone onsets, embedded images show activation patterns with variance explained, and bottom bars mark significant early- and late-change windows.

**Table S3:** Cluster-corrected activation differences. Significant intervals are reported in seconds from melody onset for the comparisons shown in Figure 3. A dash indicates that no significant cluster was detected.

| Melody | Network | M vs. early change (s) | M vs. late change (s) |
| --- | --- | --- | --- |
| M3 | 1 | 2.052–2.164 | 0.836–0.908; 2.176–2.372 |
|  | 2 | 0.484–0.744 | 0.824–1.096 |
|  | 3 | – | – |
| M5 | 1 | 0.832–0.9; 1.724–2.08 | – |
|  | 2 | 0.844–1.04; 1.672–1.84 | 1.532–1.82; 2.044–2.168 |
|  | 3 | – | – |
| M7 | 1 | 1.544–1.624; 2.388–2.524; 2.592–2.788 | 2.22–2.312; 2.384–2.504 |
|  | 2 | 1.596–1.7; 2.6–2.72 | 2.244–2.46 |
|  | 3 | 1.952–2.004; 2.616–2.68 | 2.556–2.728 |

**Figure S4:**
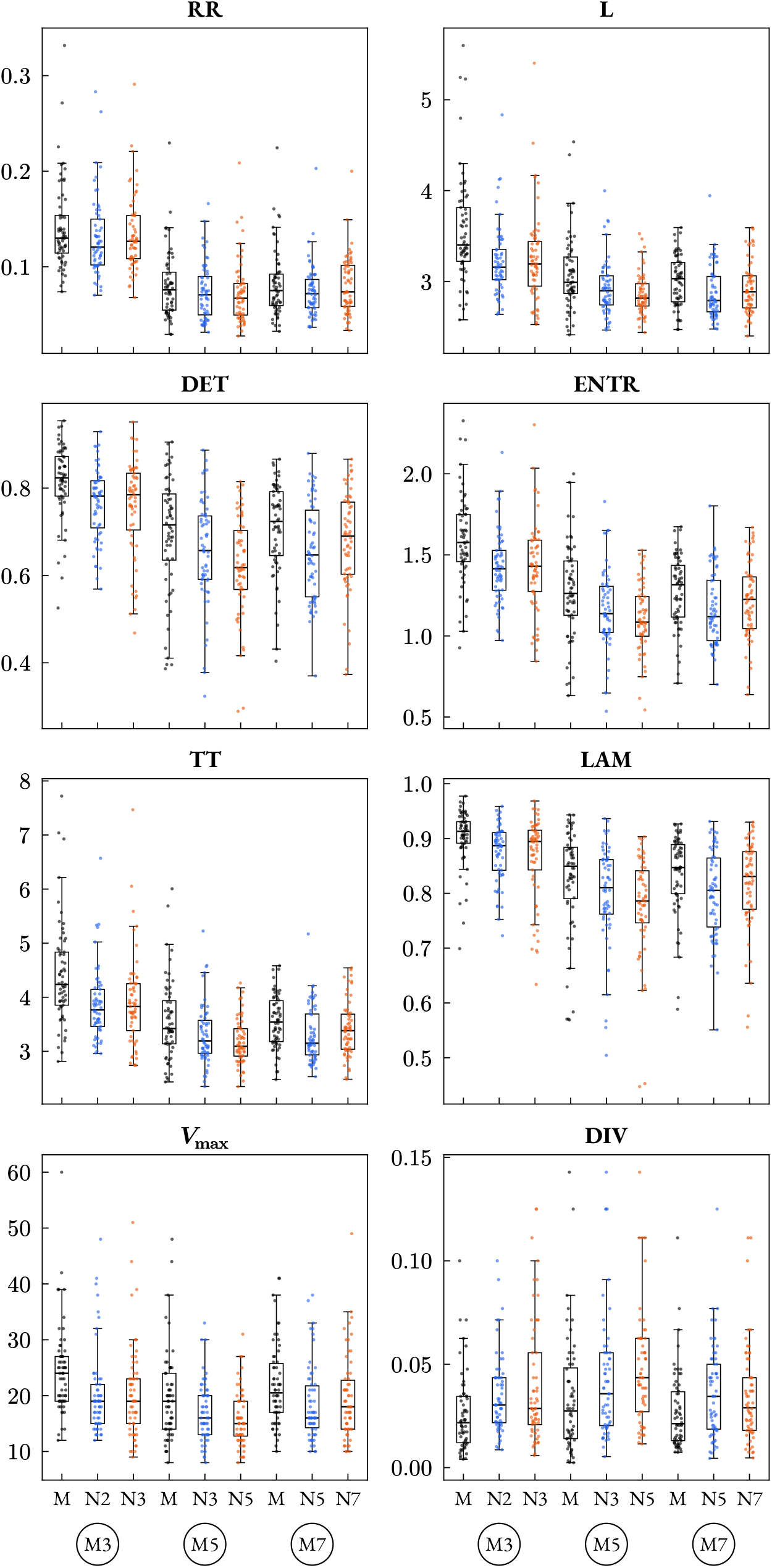
RQA metric distributions. Scatter and box plots show participant-level distributions for each RQA metric across the nine melody-length by condition-family combinations. Point color indicates condition family: 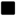 memorized (M), 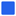 early change, and 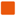 late change. N labels identify the position of the changed tone.

**Figure S5:**
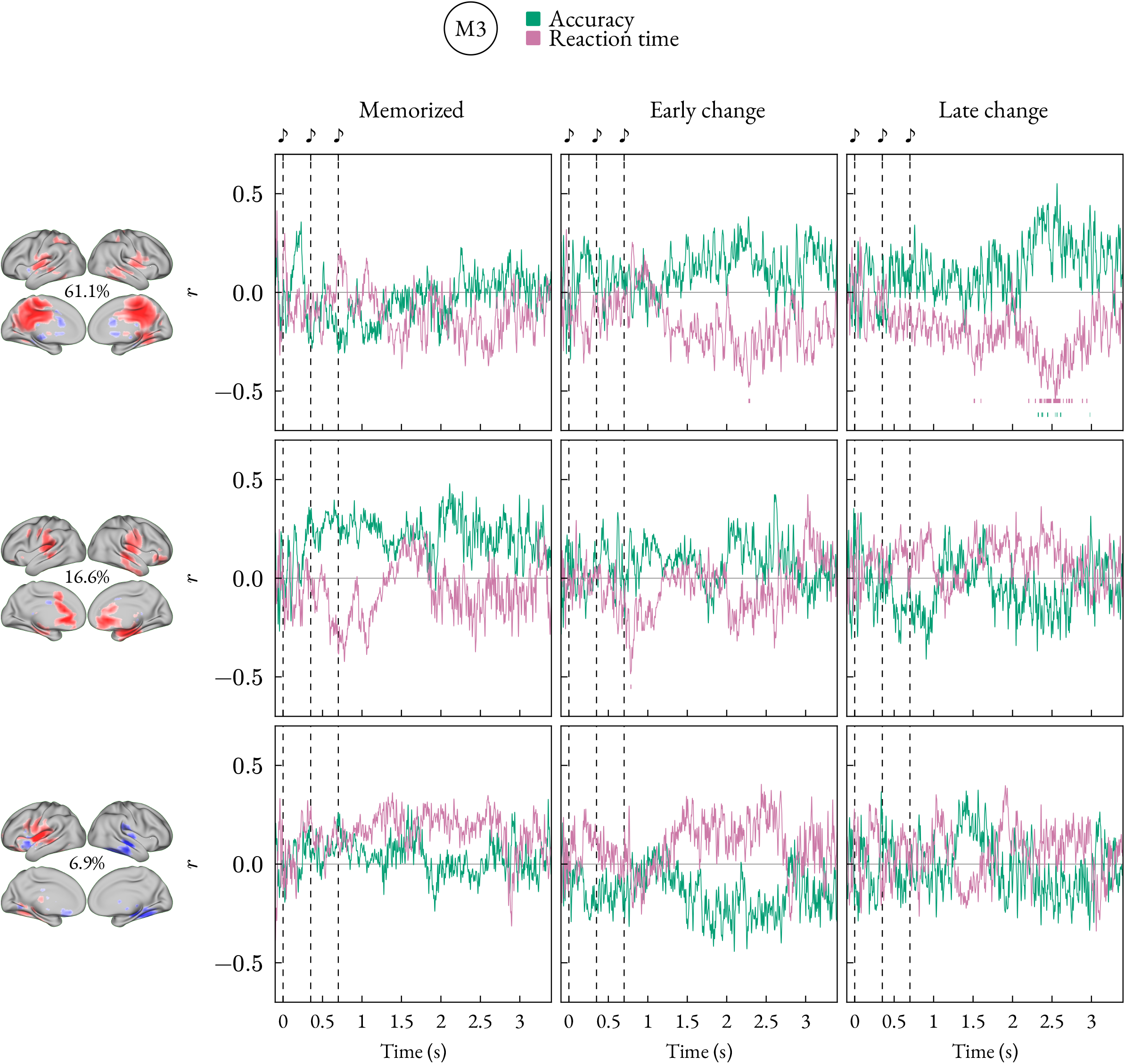
Activation-behaviour correlations. Line plots show time-resolved Pearson correlations between network activation and behavioural accuracy or mean RT for each condition family in M3. Rows show BN1–BN3, evaluated for every melody length independently of effective dimensionality, and columns show conditions. Bottom bars mark FDR-corrected significant intervals for 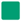 accuracy and 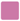 mean RT.

**Figure S6:**
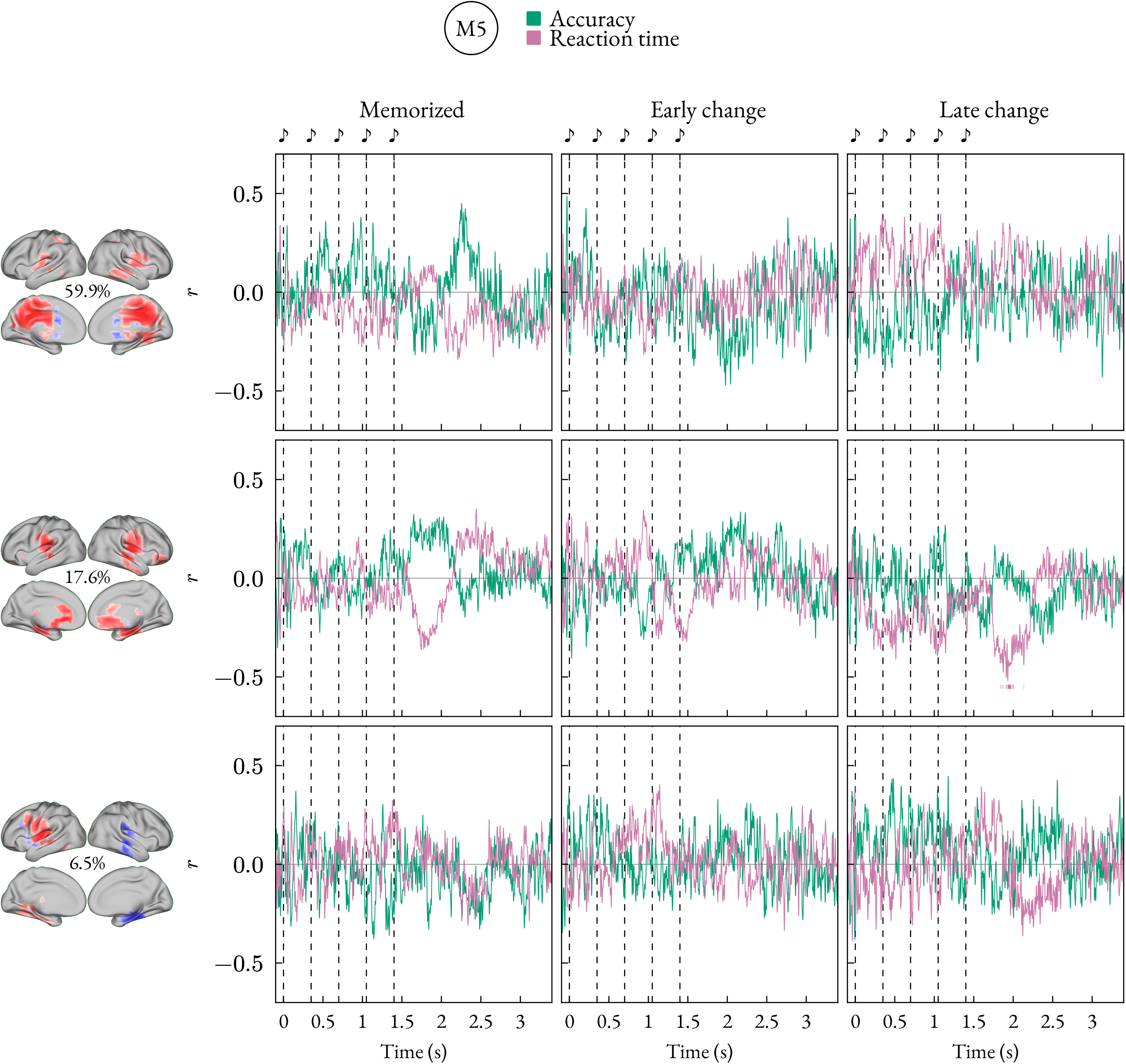
Activation-behaviour correlations for M5. Same layout as Figure S5.

**Figure S7:**
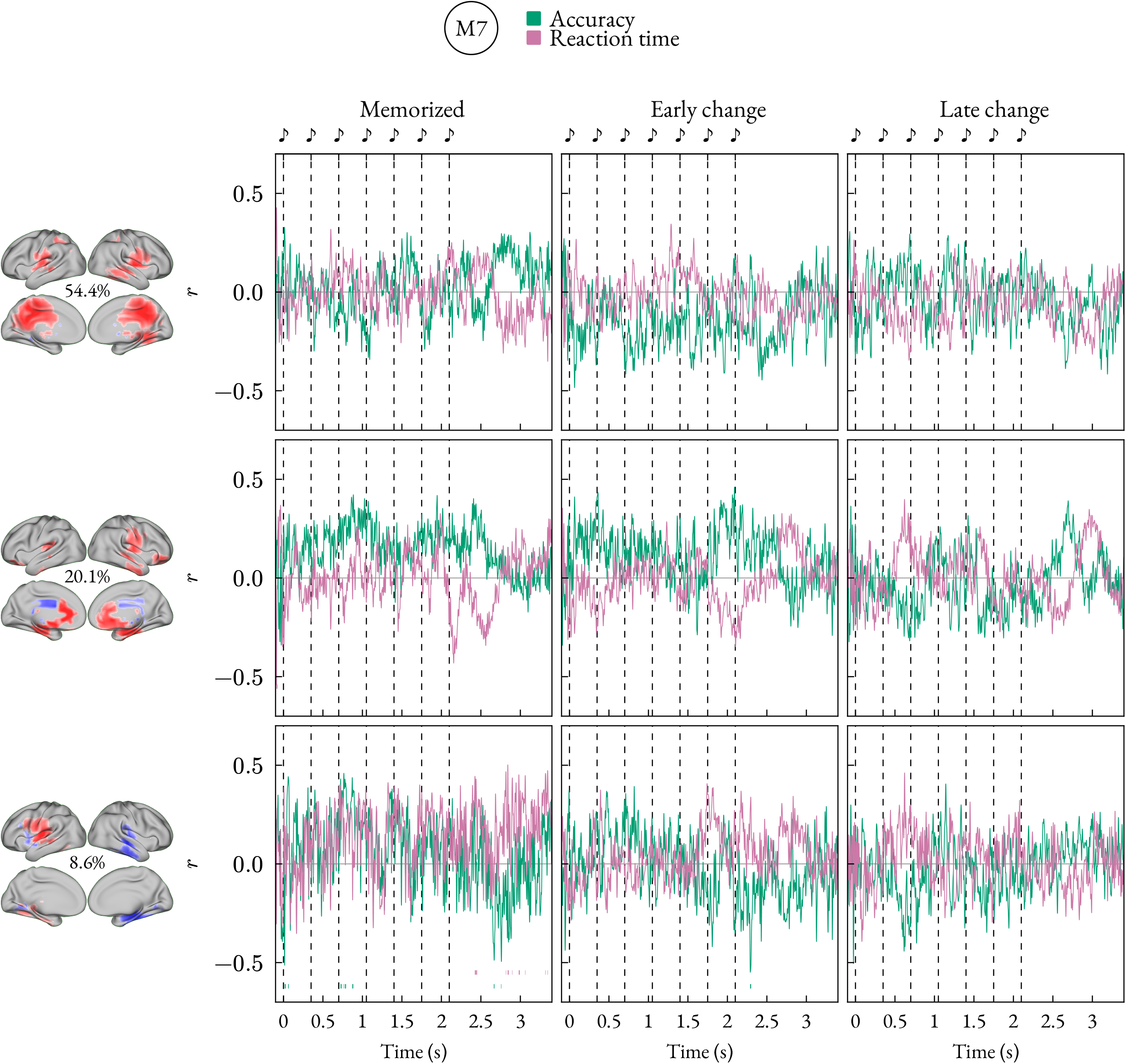
Activation-behaviour correlations for M7. Same layout as Figure S5.

**Figure S8:**
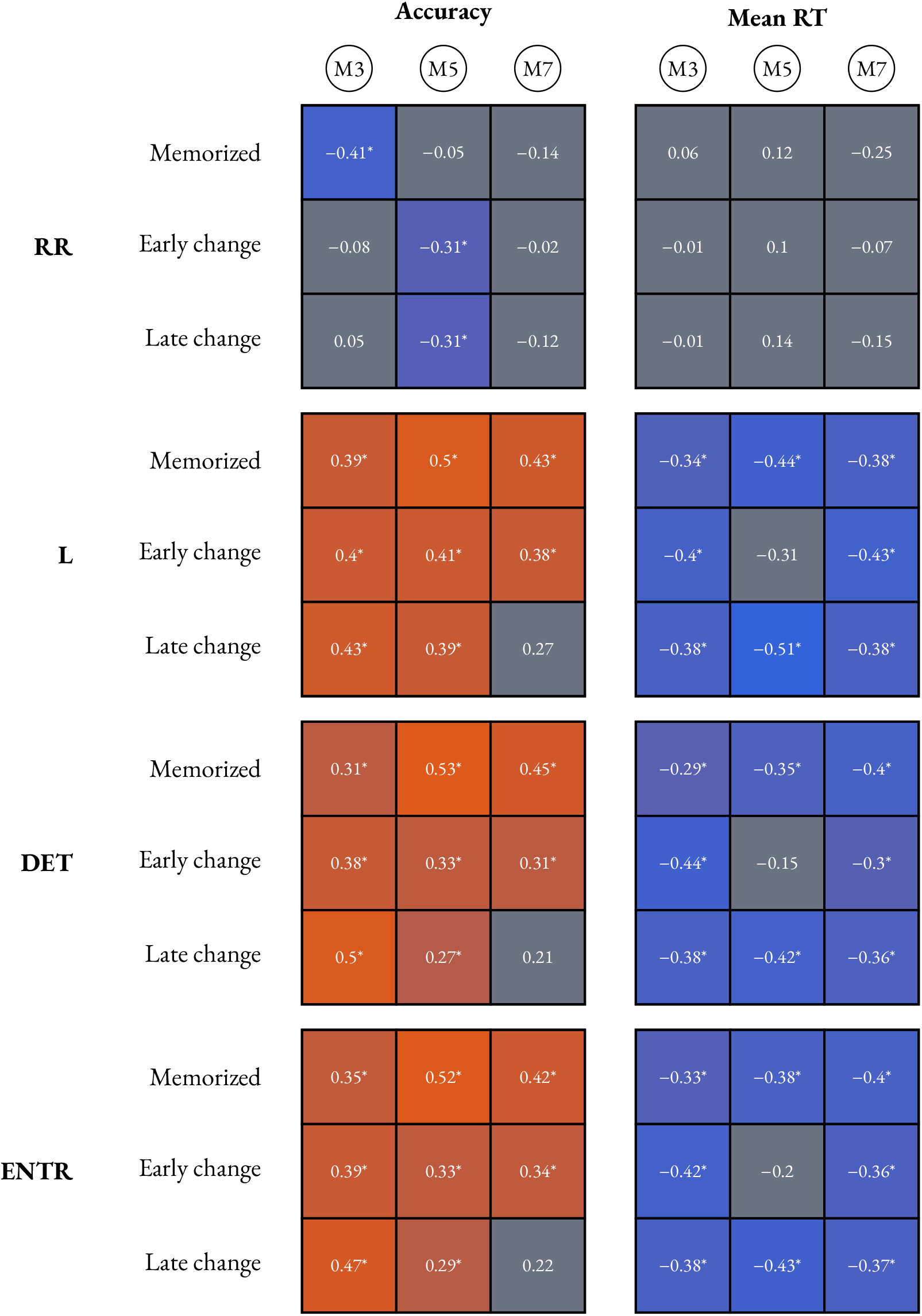
Condition-specific RQA-behaviour correlations. Each tile shows a Pearson correlation across participants between one RQA metric and one behavioural outcome for the same condition family and melody length. Behavioural values are participant-level mean correctness or mean RT for that condition, with no-response trials excluded from mean RT; RQA values are participant-level RQA summaries for the corresponding analysis window. Tile color indicates direction and magnitude, from 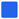 negative to 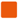 positive correlations; 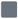 neutral tiles indicate non-significant correlations. Asterisks mark significant two-sided Pearson correlation tests, FDR-corrected across the eight RQA metrics.

**Figure S9:**
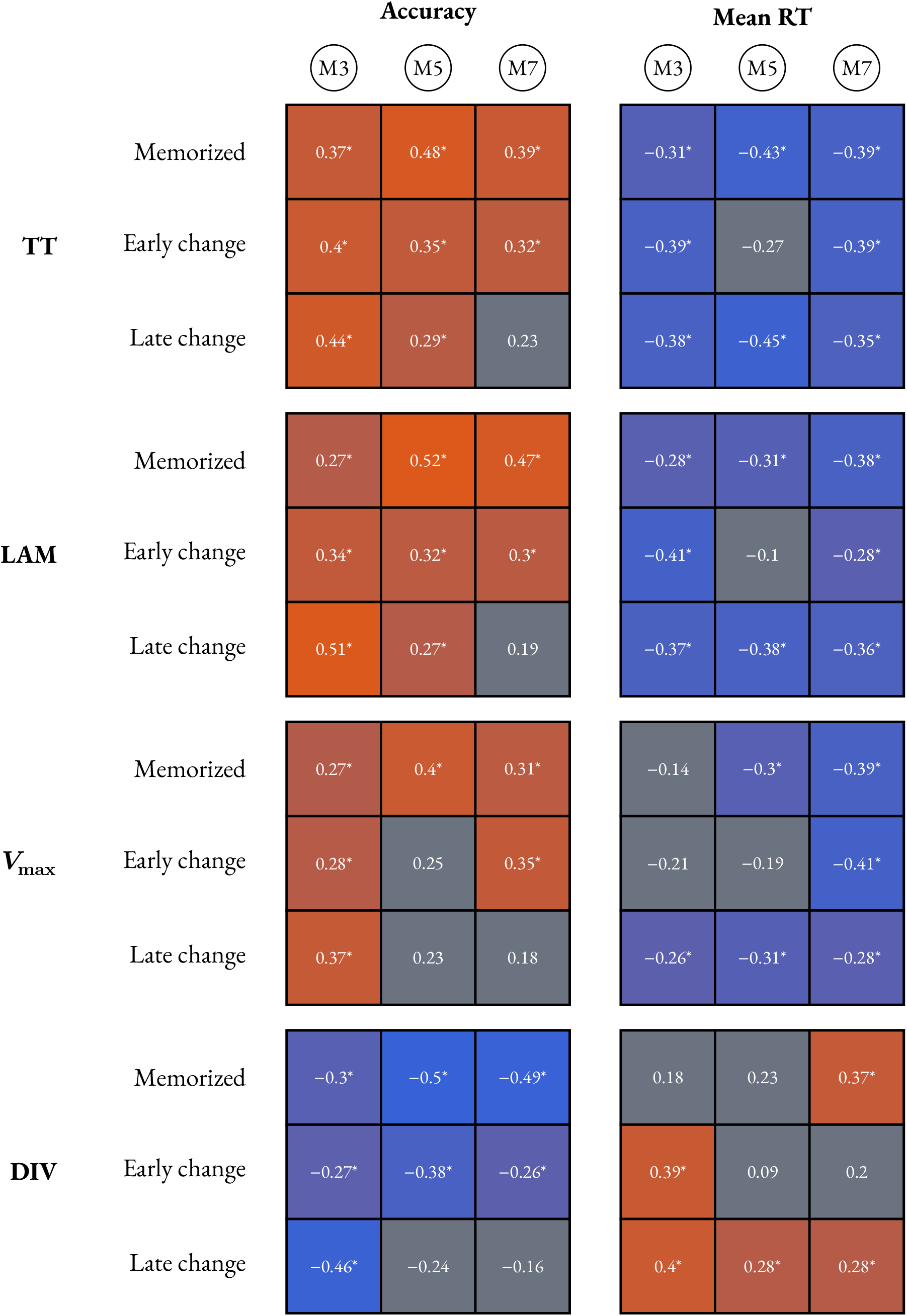
Condition-specific RQA-behaviour correlations, continued. Same layout as Figure S8.

**Figure S10:**
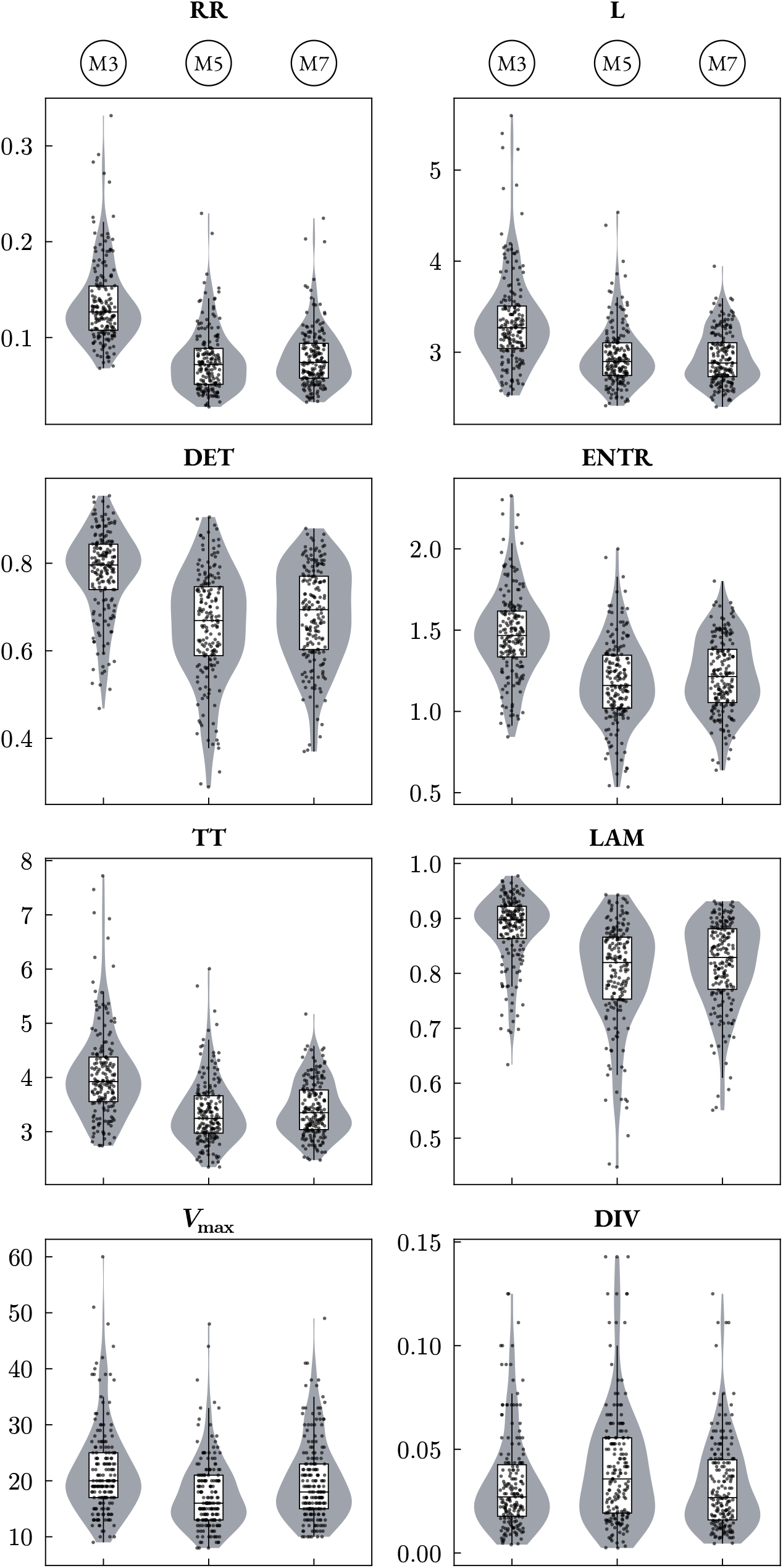
RQA metric distributions by melody length. Scatter, violin, and box plots show participant-level RQA metric distributions for M3, M5, and M7 melodies. Condition families are pooled within each melody length.

**Table S4:** Joint and metric-level contributions of RQA to accuracy beyond RMS activation amplitude. All models adjust for condition, melody length, their interaction, and RMS activation amplitude. Mixed-model rows compare this baseline with the model containing all eight standardized RQA measures; metric columns give joint-model coefficients and one-degree-of-freedom drop-one tests, with FDR correction across metrics. These individual results are descriptive because the RQA measures are collinear. Prediction rows report nested participant-wise ridge regression, in which only the RQA measures were penalized. Mean ridge coefficients summarize the 65 outer-fold fits. The confidence interval was obtained by participant-level bootstrap resampling. ΔR^2^, change in marginal R^2^.

| Statistic | Joint | RR | $L$ | DET | ENTR | TT | LAM | $V_{\max}$ | DIV |
| --- | --- | --- | --- | --- | --- | --- | --- | --- | --- |
| Standardized $\beta$ | – | –0.091 | 0.065 | 0.107 | 0.028 | –0.03 | –0.061 | 0.02 | –0.003 |
| 95% CI | – | –0.12––0.07 | –0.06–0.19 | –0.04–0.25 | –0.12–0.17 | –0.15–0.09 | –0.17–0.05 | 0–0.04 | –0.03–0.03 |
| $\chi^2$ (df) | 154.49 (8) | 46.87 (1) | 1.09 (1) | 2.03 (1) | 0.14 (1) | 0.23 (1) | 1.2 (1) | 3.24 (1) | 0.06 (1) |
| $p$ / FDR $q$ | $2.3 \times 10^{-29}$ | $6.1 \times 10^{-11}$ | 0.47 | 0.41 | 0.81 | 0.81 | 0.47 | 0.29 | 0.81 |
| $\Delta R^2$ | 0.273 | 0.091 | 0.000 | 0.002 | 0.001 | 0.000 | 0.001 | 0.002 | 0.000 |
| Mean ridge $\beta$ | – | –0.049 | 0.028 | 0.014 | 0.017 | 0.017 | 0.009 | 0.018 | –0.006 |
| Nested ridge RMSE | 0.166 | – | – | – | – | – | – | – | – |
| RMSE improvement | 11.81% | – | – | – | – | – | – | – | – |
| Participant-bootstrap<br>95% CI | 4.01%–17.65% | – | – | – | – | – | – | – | – |

**Table S5:**
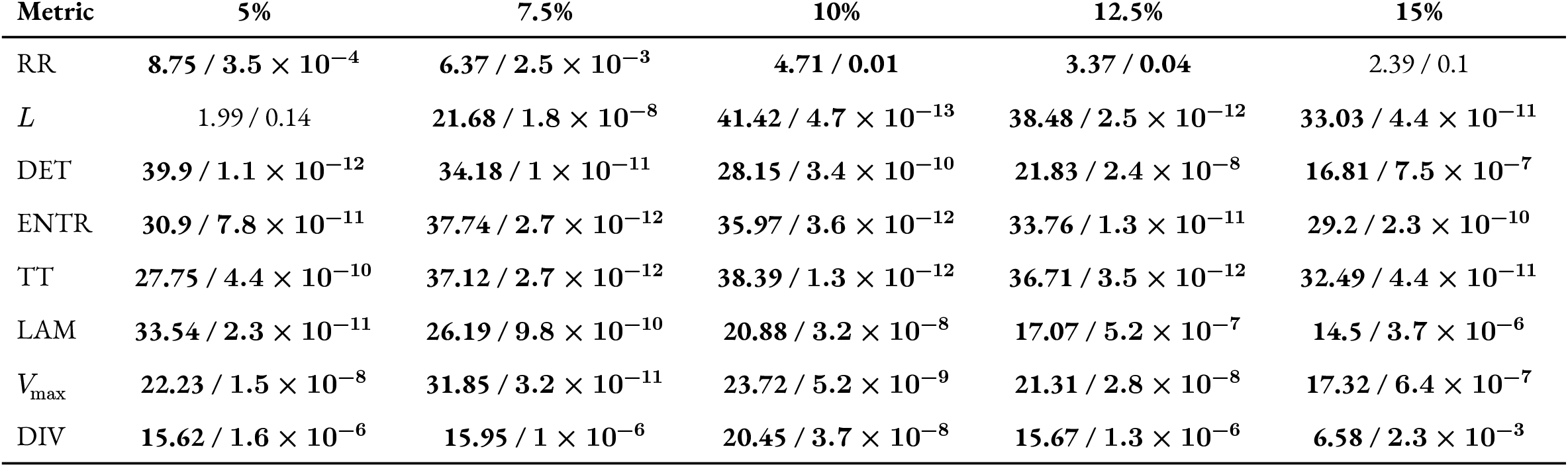
Sensitivity of RQA condition effects to the recurrence threshold. Cells report the omnibus three-level condition-effect *F* statistic followed by its FDR-adjusted *q* value. Effects were estimated across all melody lengths in the 3 × 3 repeated-measures ANOVA. FDR correction was applied across the eight metrics separately at each threshold; significant effects are bold. The 10% threshold was primary.

| Metric | 5% | 7.5% | 10% | 12.5% | 15% |
| --- | --- | --- | --- | --- | --- |
| RR | **8.75** / $3.5 \times 10^{-4}$ | **6.37** / $2.5 \times 10^{-3}$ | **4.71** / **0.01** | **3.37** / **0.04** | 2.39 / 0.1 |
| $L$ | 1.99 / 0.14 | **21.68** / $1.8 \times 10^{-8}$ | **41.42** / $4.7 \times 10^{-13}$ | **38.48** / $2.5 \times 10^{-12}$ | **33.03** / $4.4 \times 10^{-11}$ |
| DET | **39.9** / $1.1 \times 10^{-12}$ | **34.18** / $1 \times 10^{-11}$ | **28.15** / $3.4 \times 10^{-10}$ | **21.83** / $2.4 \times 10^{-8}$ | **16.81** / $7.5 \times 10^{-7}$ |
| ENTR | **30.9** / $7.8 \times 10^{-11}$ | **37.74** / $2.7 \times 10^{-12}$ | **35.97** / $3.6 \times 10^{-12}$ | **33.76** / $1.3 \times 10^{-11}$ | **29.2** / $2.3 \times 10^{-10}$ |
| TT | **27.75** / $4.4 \times 10^{-10}$ | **37.12** / $2.7 \times 10^{-12}$ | **38.39** / $1.3 \times 10^{-12}$ | **36.71** / $3.5 \times 10^{-12}$ | **32.49** / $4.4 \times 10^{-11}$ |
| LAM | **33.54** / $2.3 \times 10^{-11}$ | **26.19** / $9.8 \times 10^{-10}$ | **20.88** / $3.2 \times 10^{-8}$ | **17.07** / $5.2 \times 10^{-7}$ | **14.5** / $3.7 \times 10^{-6}$ |
| $V_{\max}$ | **22.23** / $1.5 \times 10^{-8}$ | **31.85** / $3.2 \times 10^{-11}$ | **23.72** / $5.2 \times 10^{-9}$ | **21.31** / $2.8 \times 10^{-8}$ | **17.32** / $6.4 \times 10^{-7}$ |
| DIV | **15.62** / $1.6 \times 10^{-6}$ | **15.95** / $1 \times 10^{-6}$ | **20.45** / $3.7 \times 10^{-8}$ | **15.67** / $1.3 \times 10^{-6}$ | **6.58** / $2.3 \times 10^{-3}$ |

## Notes

### Competing Interest Statement

The authors have declared no competing interest.

